# The molecular impact of cigarette smoking resembles aging across tissues

**DOI:** 10.1101/2024.03.14.585016

**Authors:** Jose Miguel Ramirez, Rogério Ribeiro, Oleksandra Soldatkina, Athos Moraes, Raquel García-Pérez, Pedro G. Ferreira, Marta Melé

## Abstract

Tobacco smoke is the main cause of preventable mortality worldwide. Smoking increases the risk of developing many diseases and has been proposed as an aging accelerator. Yet, the molecular mechanisms driving smoking-related health decline and aging acceleration in most tissues remain unexplored. Here, we characterize gene expression, alternative splicing, DNA methylation and histological alterations induced by cigarette smoking across human tissues. We show that smoking impacts tissue architecture and triggers systemic inflammation. We find that in many tissues, the effects of smoking significantly overlap those of aging in the same direction. Specifically, both age and smoking upregulate inflammatory genes and drive hypomethylation at enhancers. In addition, we observe widespread smoking-driven hypermethylation at target regions of the Polycomb repressive complex, which is a well-known aging effect. Smoking-induced epigenetic changes overlap causal aging CpGs, suggesting that these methylation changes may directly mediate aging acceleration observed in smokers. Finally, we find that smoking effects that are shared with aging are more persistent over time. Overall, our multi-tissue and multi-omic analysis of the effects of cigarette smoking provides an extensive characterization of the impact of tobacco smoke across tissues and unravels the molecular mechanisms driving smoking-induced tissue homeostasis decline and aging acceleration.

## Introduction

Tobacco smoking causes 8 million deaths annually, representing the primary cause of preventable mortality worldwide ^1^. Despite efforts to reduce smoking prevalence, the total number of smokers continues to increase, surpassing 1 billion regular smokers worldwide ^1^. Higher mortality in smokers is driven by a 3-fold increased risk of disease, including respiratory, cardiovascular, metabolic, autoimmune, renal, infectious diseases and several cancer types ^2,3^. Given its association with age-related diseases, smoking has been suggested to be an aging accelerator ^4,5^. Indeed, cigarette smoking has been associated with premature skin aging ^6^, accelerated lung function decline ^7^, telomere attrition ^8^, oxidative stress ^9^, and inflammation ^9^. Importantly, the increased disease risk does not completely reverse after smoking cessation ^10^ as 15% of deaths attributable to tobacco smoking occur in ex-smokers ^1^. In fact, recent work has shown that smoking triggers enduring alterations in the immune system, particularly with effects in adaptive responses persisting years after cessation ^11^.

Previous transcriptome and epigenome-wide studies addressing the molecular effects of smoking and the interplay with aging have been restricted to few tissues, mostly airways and whole blood. Transcriptomic studies have identified expression changes associated with both smoking and aging in human airways ^12^ and murine lungs ^13^. DNA methylation clocks that are used to measure the accumulation of DNA methylation changes with age have shown that smoking might accelerate epigenetic aging in lung and whole blood ^14–16^. Nevertheless, cigarette smoking effects are not restricted to lung or whole blood ^17,18^. Many tissues, including both directly and indirectly exposed tissues to smoke, have higher numbers of DNA adducts ^19^ and DNA mutations ^20^ as well as markers of oxidative damage and inflammation ^9^. In addition, smoking-associated DNA methylation changes in lung, colon and adipose tissues have been shown to mediate gene expression changes that could impact lung function and metabolic health ^17,18^. Finally, smoking-associated diseases involve the cardiovascular, metabolic and immune systems ^3^, providing further evidence that the impact of cigarette smoking is systemic. However, what molecular and histological consequences of smoking could lead to aging acceleration in human tissues remains unexplored.

Here, we use data from the Genotype-Tissue Expression (GTEx) project ^21^ to systematically analyze the transcriptomic and histological impact of cigarette smoking across dozens of human tissues previously unexplored. We identify smoking-induced differences in gene expression, alternative splicing, DNA methylation and tissue architecture across tissues and identify those effects that are simultaneously associated with both smoking and aging. Using ex-smoker information, we classify smoking-associated changes as reversible or non-reversible and address whether smoking effects overlapping those of aging are more persistent. Overall, our multi-tissue and multi-omic analysis provides a comprehensive overview of the systemic and tissue-specific impact of cigarette smoking and how these may drive accelerated tissue aging. Our study sheds light on the long-term health effects and associated risks of smoking, as well as the potential health benefits of smoking cessation.

## Results

### Cigarette smoking impacts gene expression and alternative splicing in most human tissues

The transcriptomic impact of cigarette smoking has been studied in a limited number of tissues^12,22^. To perform a systematic quantification of the effect of cigarette smoking on gene expression across human tissues, we used the Genotype Tissue Expression (GTEx) v8 data release ^21^ (Fig. 1). We selected 46 tissues with at least 80 samples comprising 12,654 RNA-sequencing samples from 717 donors, including 316 smokers, 253 never smokers and 148 ex-smokers. We performed differential gene expression analysis per tissue comparing smokers and never smokers and controlling for technical and biological covariates (Methods, Table S1). The number of smoking-associated differentially expressed genes (smoking-DEGs) varied across tissues (FDR < 0.05, Fig. 2a, Fig. S1a, Table S2). Lung (2,656) followed by thyroid (1,345) and esophagus mucosa (1,096) had the largest number of smoking-DEGs. Downsampling to an equal number of samples per tissue showed consistent results, with lung, pancreas, thyroid, and esophagus mucosa having the largest number of smoking-DEGs (Fig. 2b, Fig. S1b). Importantly, we were able to reproduce previous findings as the smoking-DEGs here identified significantly overlap with those identified in previous studies for the same tissue (two-tailed Fisher’s exact test; p-value < 0.05) (Fig. S1e)^22–27^.

**Fig. 1.**
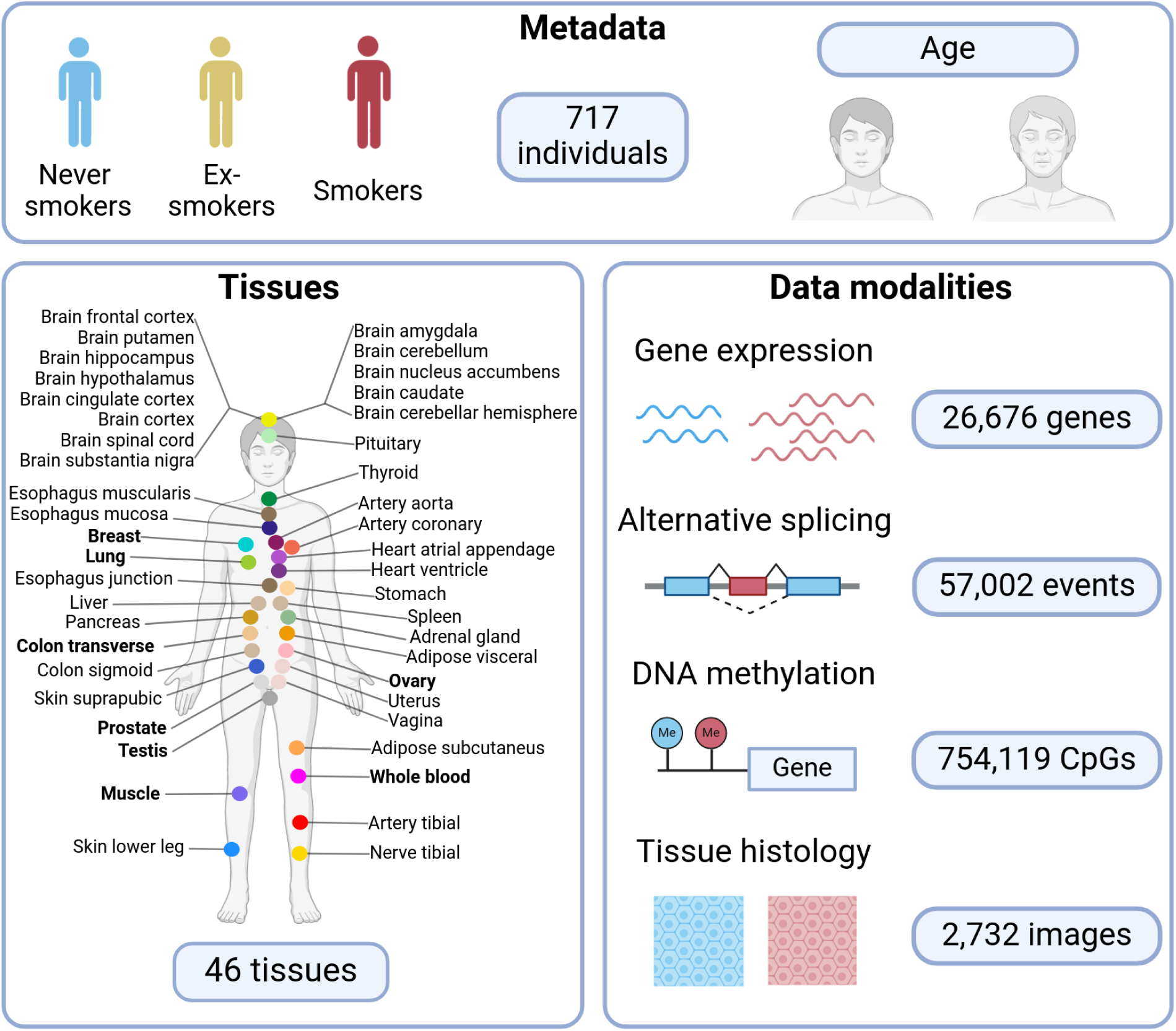
Individuals, tissues and data modalities analyzed. For all tissues, RNA-sequencing and histological images are available. Tissues in bold additionally have DNA methylation information. We classified GTEx individuals according to their smoking status.

**Fig. 2.**
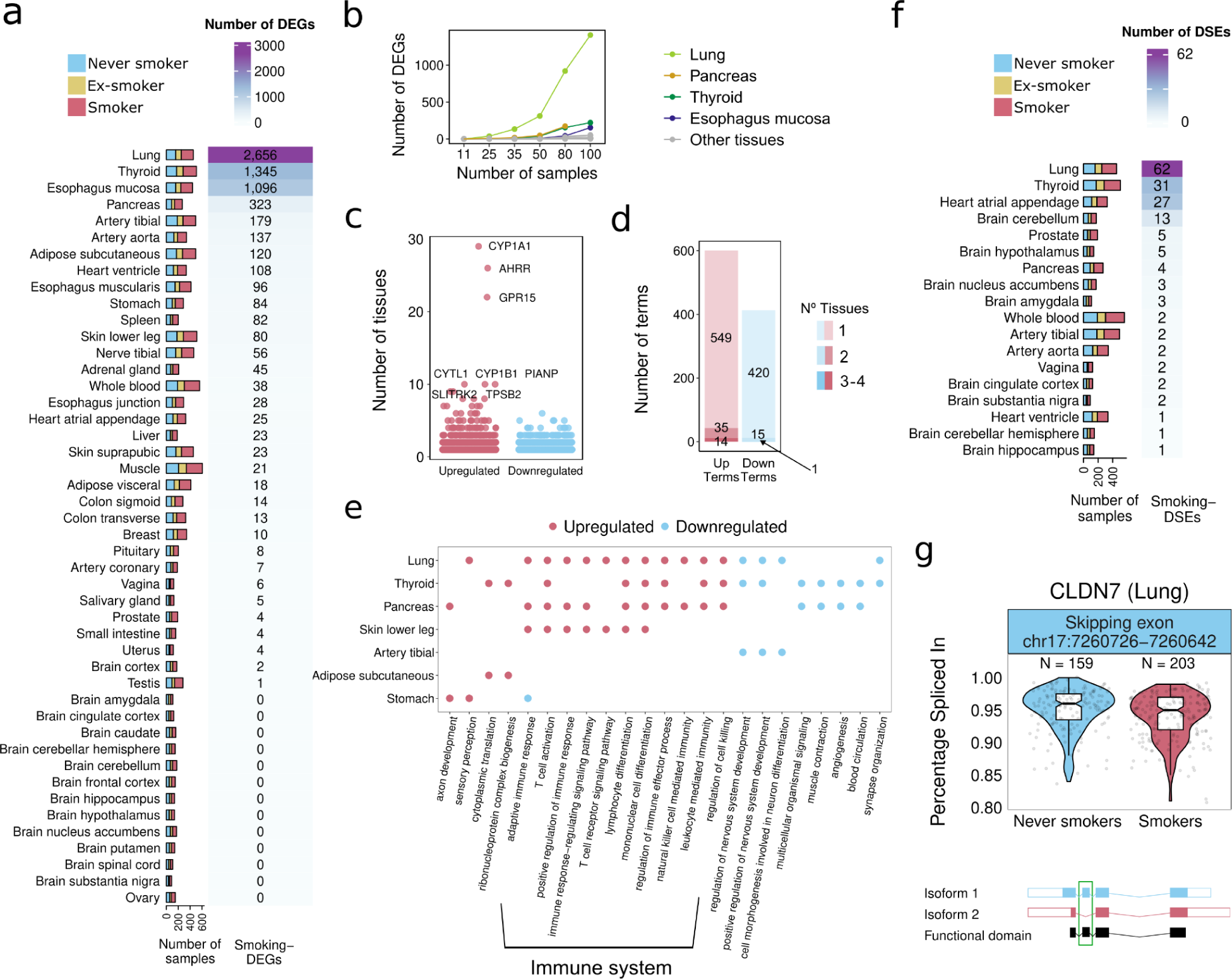
Smoking differential gene expression and alternative splicing analysis. **a**, Tissue sample size (left) and number of smoking-DEGs per tissue (right). **b**, Number of smoking-DEGs per tissue at varying sample sizes while keeping an equal number of smokers and never smokers. **c**, Number of tissues in which the smoking-DEGs are differentially expressed. Gene names are shown for smoking-DEGs in more than 9 tissues. **d**, Number of tissue-specific and tissue-shared gene ontology enriched terms (FDR < 0.05, minimum gene count > 5). **e**, Most enriched and summarized gene ontology terms (biological processes) for smoking up- (red) and down-(blue) regulated genes. **f**, Tissue sample size (left) and number of smoking-DSEs per tissue (right). **g**, PSI values in smokers and never-smokers for a skipped exon in *CLDN7* (top) and visual representation of the two most expressed isoforms involved in the exon skipping event (bottom). Isoform 1 is ENST00000360325 and Isoform 2, ENST00000538261.

Smoking effects are largely tissue-specific, with 86% of smoking-DEGs altered in a single tissue (Fig. 2c). However, 8 genes were found upregulated in 9 or more tissues (Fig. 2c, Fig. S1d). Of these, AHRR, CYP1A1 and CYP1B1 are known to be upregulated upon direct exposure to polycyclic aromatic hydrocarbons (PAHs)^28^, suggesting that toxic compounds reach tissues not directly exposed to tobacco smoke. Other highly shared and upregulated genes, such as GPR15, PIANP, CYTL1 and TPSB2, are involved in immune system functions and inflammation ^29–32^,. Functional enrichment showed little overlap across tissues, indicating that most smoking-altered pathways are tissue-specific (Fig. 2d, Table S3-4). For example, cilium assembly pathways are downregulated in the lung, consistent with reported cilia dysfunction in smokers ^33^ and cancer-related pathways are downregulated in the thyroid, which is intriguing as smokers are at lower risk of developing thyroid cancer ^34^(Fig. 2e, Fig. S2). The few pathways that are shared across tissues are mostly upregulated and associated with immune function, consistent with increased inflammation in smokers ^35–37^(Fig. 2e).

Alternative splicing has been associated with tobacco smoking in blood ^38,39^. We computed the “percentage spliced-in” (PSI) for seven types of alternative splicing events ^40^ and performed differential analysis to test the association between PSI variation and smoking, while correcting for known sources of technical variation (Methods). We found differentially spliced events (smoking-DSE) in 17 tissues (Fig. 2f, Table S5). The most affected tissues were lung (65), thyroid (34) and heart (27). In half (48%) of the smoking-DSEs both the inclusion and exclusion of the event are associated with at least one protein-coding isoform, of which 46% are associated with direct changes in annotated protein domains ^41^ (Methods), including the *CLDN7* case (Fig. 2g). Most of the other half of smoking-DSEs (44%) are changes between protein-coding and non-coding isoforms, with smoking being associated with the loss of coding potential (64%, binomial test; p-value = 0.015).

Overall these results show that smoking affects gene expression and alternative splicing in multiple tissues leading to both tissue-specific changes and systemic inflammation.

### Cigarette smoking induces histological changes in lung and thyroid

We hypothesized that the observed smoking-induced gene expression effects could be in part explained by histological changes in those tissues. To further investigate the impact of smoking on tissue architecture, we selected the four tissues with the largest number of smoking-DEGs (Fig. 1a lung, thyroid, pancreas and esophagus mucosa) and analyzed their histological images (Fig. 3a). We trained a convolutional neural network per tissue to classify histology images as smokers or never smokers (Fig. S3a). All classifiers have higher performance than expected by chance (Fig. 3b), suggesting that smoking has an impact on the four tissue’s histology. The lung classifier has the best performance (AUC = 0.85), consistent with the known damaging effects of smoking in this tissue ^42–45^. Visual inspection of the lung images revealed a substantial presence of macrophages (Fig. 3c). Cell type deconvolution analysis (Methods) confirmed that smokers have a higher proportion of macrophages (z-test; FDR = 2.26e-06), according to previous reports ^42–44^(Fig. 3d, Table S6).

**Fig. 3.**
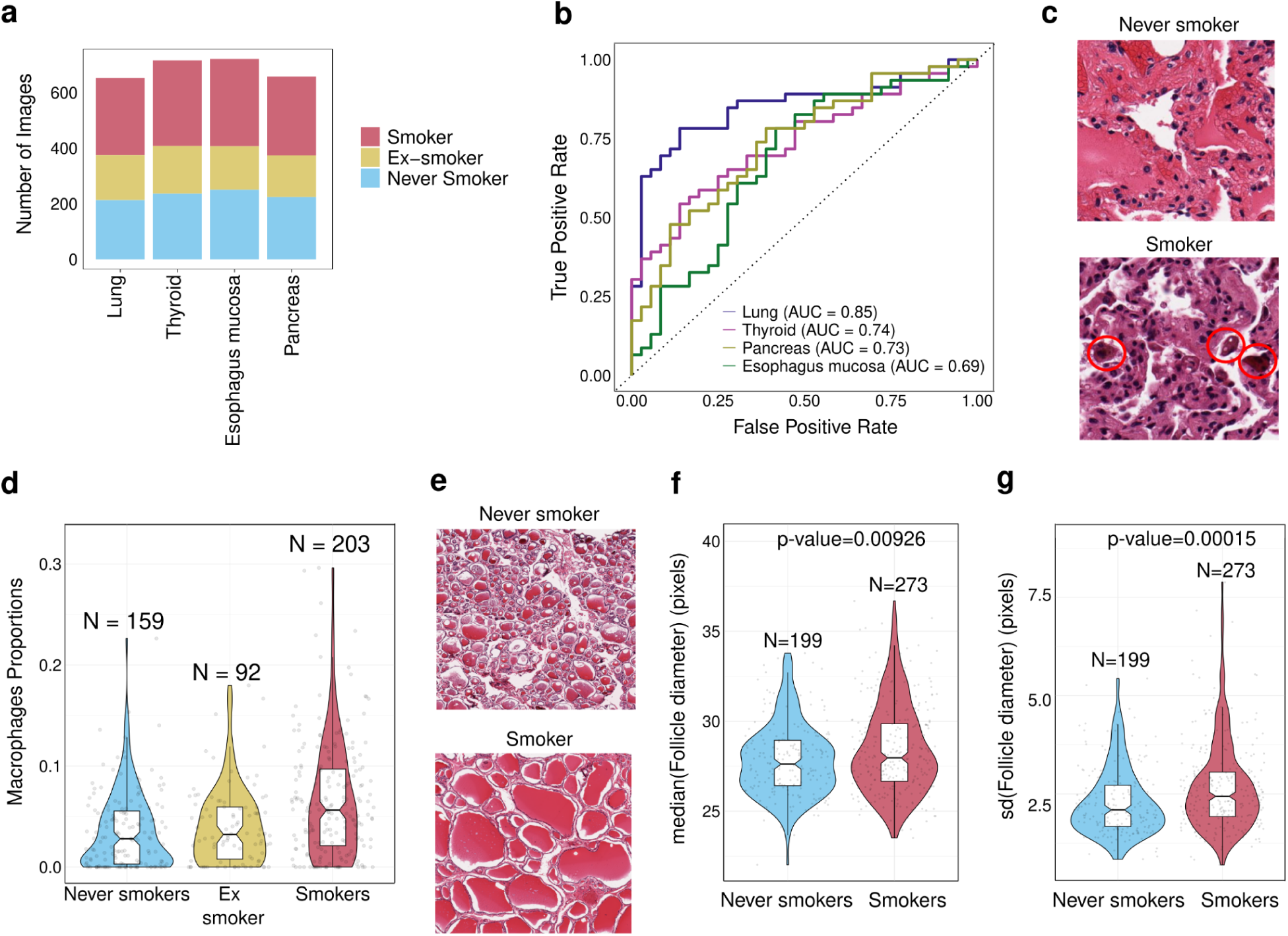
Impact of smoking on histology. **a**, Number of images analyzed per tissue and per donor smoking status. **b**, Receiver Operating Characteristic (ROC) curves of the tissue histology classifiers. **c**, Representative images of a never smoker and a smoker’s lung. Brown macrophages are highlighted in red circles. **d**, Estimated proportion of macrophages in never smokers and smokers. **e**, Representative images of a never smoker and smoker’s thyroid. Smoker thyroids have larger colloid-containing follicles. **f**, Mean (per donor) of median (per tile) diameter of thyroid follicles. p-value obtained from a Wilcoxon test. **g**, Median (per donor) of standard deviation (sd) (per tile) diameter of thyroid follicles. p-value obtained from a Wilcoxon test.

The thyroid classifier has the second-best performance (AUC=0.74), consistent with thyroid being the second tissue with the largest number of smoking-DEGs. We visually inspected thyroid images and observed that smokers have bigger colloid-containing follicles (Fig. 3e). Thyroid follicles are the storage unit of inactive thyroid hormones. We used *CellProfiler*^46^ to identify thyroid follicles automatically from the images and compute size and shape metrics (Fig. S3b, Table S7). We find that smokers have larger mean (Fig. 3f) and standard deviation (Fig. 3g) follicle diameters compared to never smokers. This is consistent with the well-known association between smoking and goiter, which is an irregular growth of the thyroid gland ^34^. Pancreas and esophagus mucosa models show similar discriminatory performance, highlighting that cells and tissues undergo structural changes as a result of smoking.

### Smoking and aging alter gene expression across tissues in the same direction

Common gene expression changes between smoking and aging in the respiratory tract have suggested that smoking could be an aging accelerator ^12,13^. To assess if this extends to other tissues, we tested if the overlap between smoking-DEGs and age-DEGs is higher than expected by chance. For comparison, we performed the same test on three other demographic traits: sex, ancestry and Body Mass Index (BMI), which are also known to influence gene expression variation ^47^. Age is the demographic trait with the largest number of tissues with significant overlap with smoking-DEGs (n=8)(Fig. 4a-b) (one-tailed Fisher’s exact test; FDR < 0.05). For the 6 tissues with more than 10 smoking-age-DEGs, 5 have a significant bias in the direction of change (chi-squared test; FDR < 0.05). In all cases, the direction of the effect is concordant (Fig. 4c, Fig. S4a-c). Among the concordant smoking-age-DEGs, we find pathways significantly enriched in four tissues with the strongest associations related to immune functions for upregulated genes in pancreas and skin (Table S8).

**Fig. 4.**
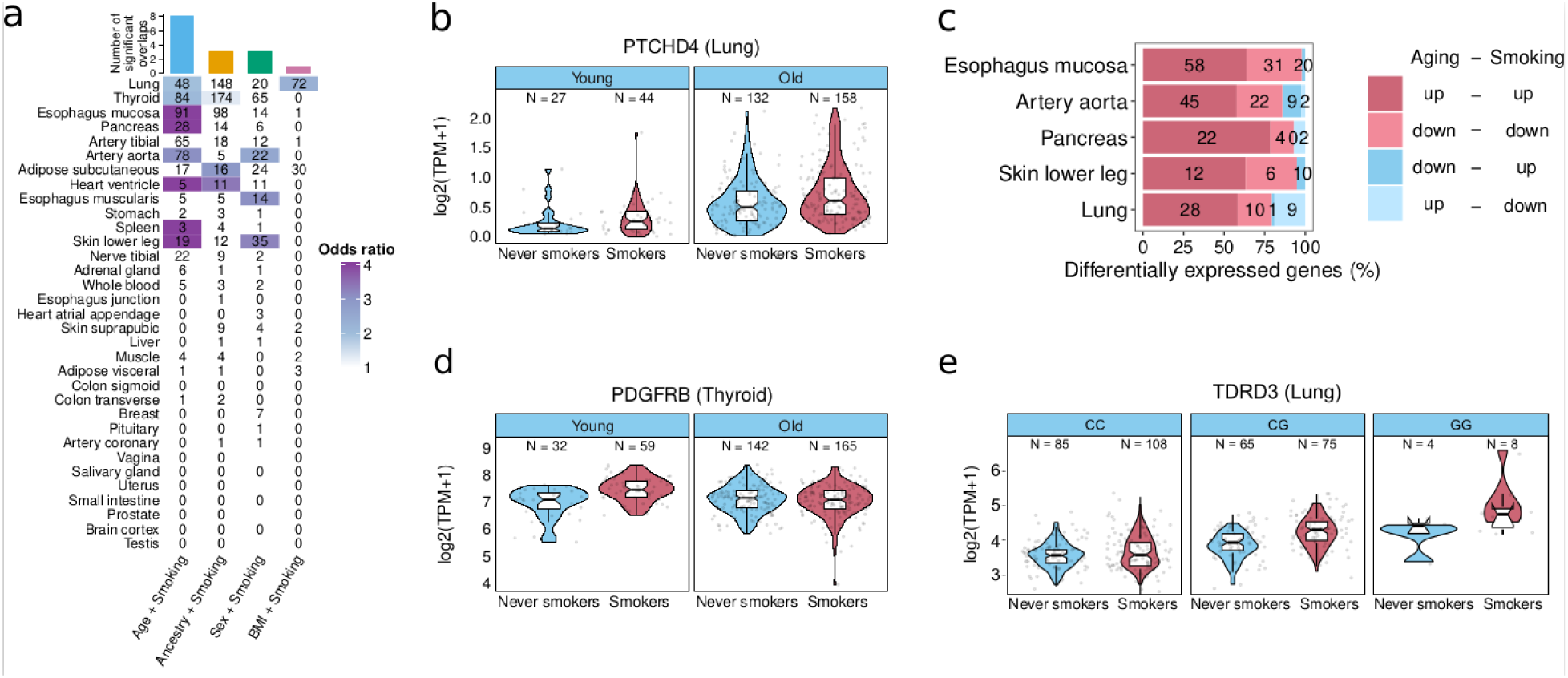
Impact of smoking and aging on gene expression. **a**, Number of smoking-DEGs that are also DE with one demographic trait per tissue. Coloured cells indicate higher-than-expected overlaps (Fisher’s exact tests; FDR < 0.05). Tissues are sorted from highest (top) to lowest (bottom) sample size. **b**, Example of a gene with additive effects for smoking and age. **c**, Tissues with significant concordance in the direction of change (up- or down-regulation) for smoking-age-DEGs across tissues (chi-squared tests; FDR<0.05). Tissues are sorted from lowest (top) to highest (bottom) p-value. **d**, Example of a gene with an interaction effect between smoking and age. **e**, Expression of *TDRD3* stratified by genotype at rs7924558 position and smoking status.

We then tested if the effects of smoking differed depending on intrinsic donor characteristics such as age or genetic background. First, we performed differential gene expression analysis extending the previous models adding an interaction term for smoking and each of the four traits (age, sex, genetic ancestry and BMI). We identified 10 genes with significant interactions (Fig. S4d). For instance, *PDGFRB,* a gene involved in premature aging ^48^, is upregulated in young smokers (Fig. 4d). Then, to test whether the impact of smoking on gene expression varied depending on *cis*-genetic effects, we performed expression quantitative trait loci analysis (eQTL) across tissues including a genotype x smoking interaction term (Methods). We identified one significant interaction in *TDRD3*, a chromatin remodeler gene, at position *rs7924558* in lung (Fig. 4e).

Overall, we find that smoking acts additively with aging changing expression across tissues in the same direction, while interactions are rare, supporting the idea that smoking acts as a systemic tissue aging accelerator, often by increasing inflammation.

### Smoking induces hypermethylation at targets of the Polycomb repressive complex

A recent study generated DNA methylation data for 9 GTEx tissues and showed that smoking is associated with DNA methylation changes in the lung and colon ^18^. We wanted to address if smoking-induced DNA methylation were similar to age-associated methylation changes, similarly to what we have observed in gene expression in the same cohort. To address this, we performed differential methylation analysis with smoking correcting for PEER values across the 9 tissues (Fig. S5a) (Methods). We identified 96,826 differentially methylated positions (smoking-DMPs) in the lung and 87 in the colon transverse (Fig. 5a, Table S9)^18^. Smoking-DMPs significantly overlap those identified in previous studies (Fig. S5b; Fisher’s exact tests; FDR < 0.05)^18,27,49–53^. However, we identify a larger number of smoking-DMPs in the lung and a higher proportion of hypermethylation compared to other studies ^18,27,49,50,52–54^. This higher proportion of hypermethylation might be due to our analysis detecting DMPs with smaller effect sizes. Indeed, hypomethylated CpGs have larger effect sizes (Fig. S5c) and when downsampling to smaller sample sizes, we retrieve higher proportions of hypomethylation (Fig. S5d), suggesting that they are easier to detect.

**Fig. 5:**
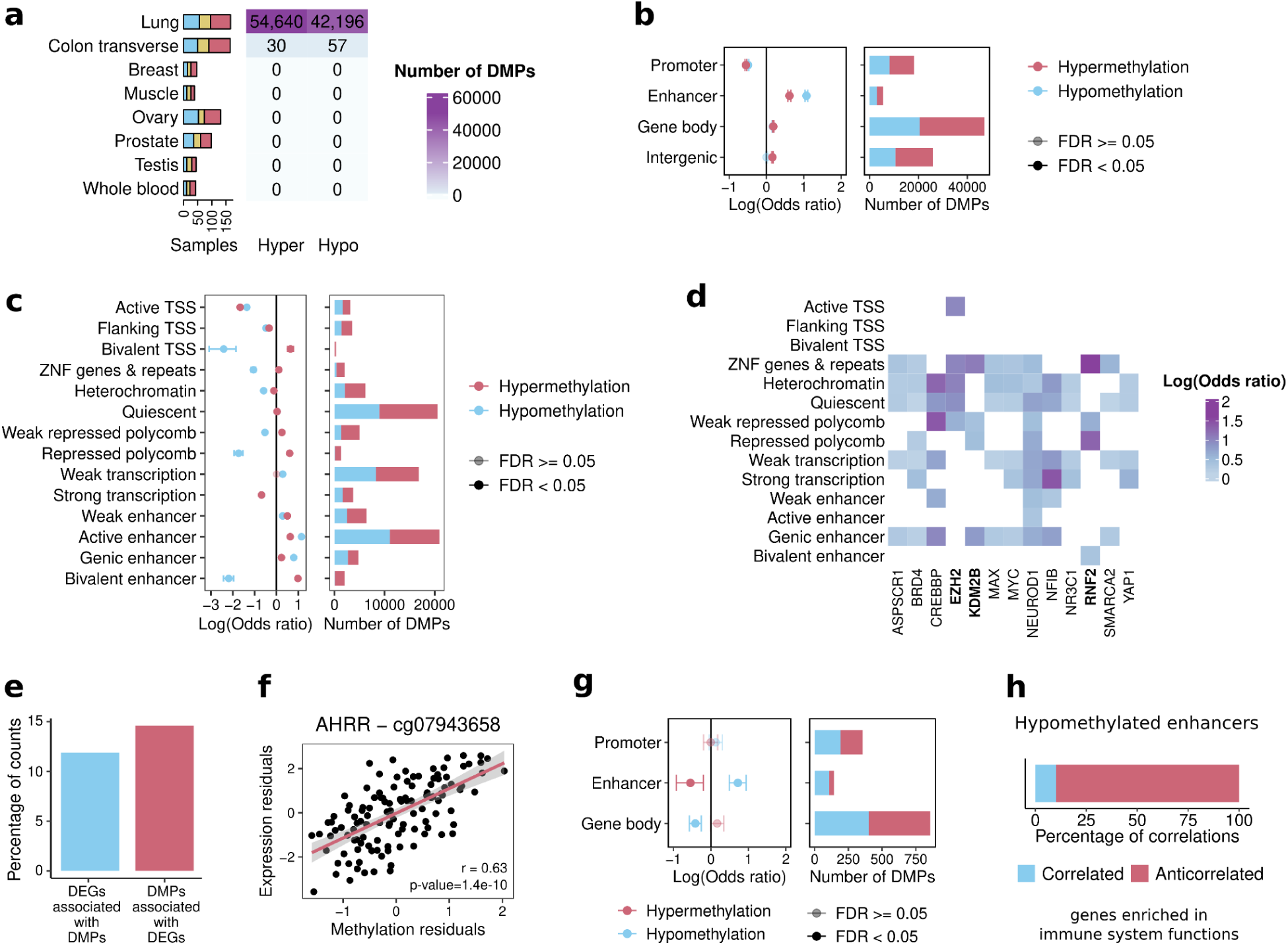
Association of smoking with DNA methylation. **a**, Number of samples (left) and smoking-DMPs (right) across tissues. **b**, Smoking-DMPs enrichment at regulatory regions. **c**, Smoking-DMPs enrichment at chromatin states **d**, TFBSs enriched in hypermethylated smoking-DMPs in more than 3 chromatin states. **e**, Percentage of smoking-DEGs associated with at least one smoking-DMP and vice versa. **f**, Correlation of methylation residuals in cg25648203 with expression residuals of AHRR. **g**, Enrichment of significantly correlated DMPs-DEGs at regulatory regions. **h**, Percentage of positive and negative correlations for hypomethylated enhancers (top), and pathway in which these correlations are enrichment (bottom).

We then characterized our newly identified smoking-DMPs according to genomic context because DNA methylation can have distinct regulatory effects when occurring in different genomic regions ^55^. In lung, smoking-DMPs are depleted in promoters and CpG islands and enriched in enhancers and open sea regions (Fig. 5b, Fig. S5e-f). Using lung-specific chromatin segmentations as defined by ChromHMM ^56^, we find bivalent enhancers, bivalent TSS, and Polycomb-repressed regions enriched in hypermethylated smoking-DMPs and depleted in hypomethylated smoking-DMPs (Fig. 5c). When looking at genes having smoking-DMPs nearby, we observe these genes are mostly associated with developmental functions, with the strongest signal in hypermethylated smoking-DMPs at active enhancers, active and flanking TSS, and quiescent regions (Table S10). Whereas enrichment of smoking-DMPs at enhancers has been previously observed ^49,53^, the observation that hypermethylated smoking-DMPs are enriched at bivalent and Polycomb-repressed regions is novel. These regions have in common low basal levels of methylation (Fig. S6a) and are marked by the repressive histone mark H3K27me3 placed by the Polycomb repressive complex ^57^. In addition, hypermethylation at Polycomb targets has been previously reported as a systemic effect of aging ^58^ . To check if this effect extends to other tissues, we looked at the location of DMPs reported in a whole blood study with a larger sample size ^53^. Hypermethylated DMPs are significantly enriched in bivalent TSS (Fig. S6b) (Fisher’s exact test; OR=4, FDR=0.029). The second highest odds ratio, although marginally not significant, was at bivalent enhancers (Fisher’s exact test; OR=2.3, FDR=0.06). This suggests that smoking-induced hypermethylation at bivalent and Polycomb-repressed regions is shared across tissues. Consistent with this, among the TFs enriched in hypermethylated smoking-DMPs shared across most chromatin regions (Fig. 5d, S6d) are TFs part of the Polycomb repressive complex (EZH2, RNF2 and KDM2B).

We then wanted to explore if the observed smoking-induced methylation correlates with gene expression changes. Restricting our analysis to CpG-gene pairs described in the EPIC annotation, we observe very few (11-15%) DMP-DEG pairs (Fig. 5e-f) suggesting that, in most cases, smoking has independent effects on DNA methylation and gene expression. In addition, from the DMP-DEGs, only 12.6% of the smoking-DMPs (1,356) were correlated with the associated smoking-DEG, mostly negatively (68%, Fig. S7a). Interestingly, the correlated smoking-DMPs are enriched in hypomethylated enhancers (Fig. 5g), where 90% of the correlations are negative (Fig. 5h) and at genes enriched in functions related to the immune system (Table S12), suggesting that previously observed smoking-induced DNA hypomethylation at enhancers ^49,53^ is related to immune upregulation. Interestingly, hypermethylated CpGs at bivalent and Polycomb-repressed chromatin states show fewer significant correlations with gene expression (4%) than other hypermethylated positions with similar effect sizes (15%) (binomial test; p-value=5.16e-08), suggesting that smoking-induced hypermethylation at targets of the Polycomb repressive complex does not significantly impact gene expression.

Overall, we find that smoking drives hypermethylation at target regions of the Polycomb repressive complex, although with marginal effects in gene expression.

### Smoking shows additive effects with aging at Polycomb binding sites and alters aging driver CpGs

Cigarette smoking has been associated with increased epigenetic aging as age-predictive models trained on methylation marks consistently predict smokers as older than their chronological age ^14,15^. Our analysis shows higher hypermethylation at bivalent and polycomb repressed regions (Fig. 5c), which is known to occur also in aging ^58^. Thus, we tested whether smoking and aging impact the same CpGs in the same direction across tissues. We find that smoking-DMPs significantly overlap age-DMPs in the same direction in both lung and colon (Fig. 6a) (Chi-square test; FDR<0.05). The strongest enrichment of age-smoking-DMPs occurs at hypomethylated enhancers (Fig. 6b) and open sea regions (Fig. S7b). When we focus on chromatin states, the strongest enrichments occur for hypermethylation in bivalent enhancers (25-fold), bivalent TSS (10-fold) and Polycomb-repressed regions (10-fold)(Fig. 6c), regions associated with development that are known to be hypermethylated with aging ^59,60^. Functional enrichment analysis on the genes associated with smoking-age-DMPs shows that hypermethylation in active TSS is also associated with developmental genes (Table S10).

**Fig. 6.**
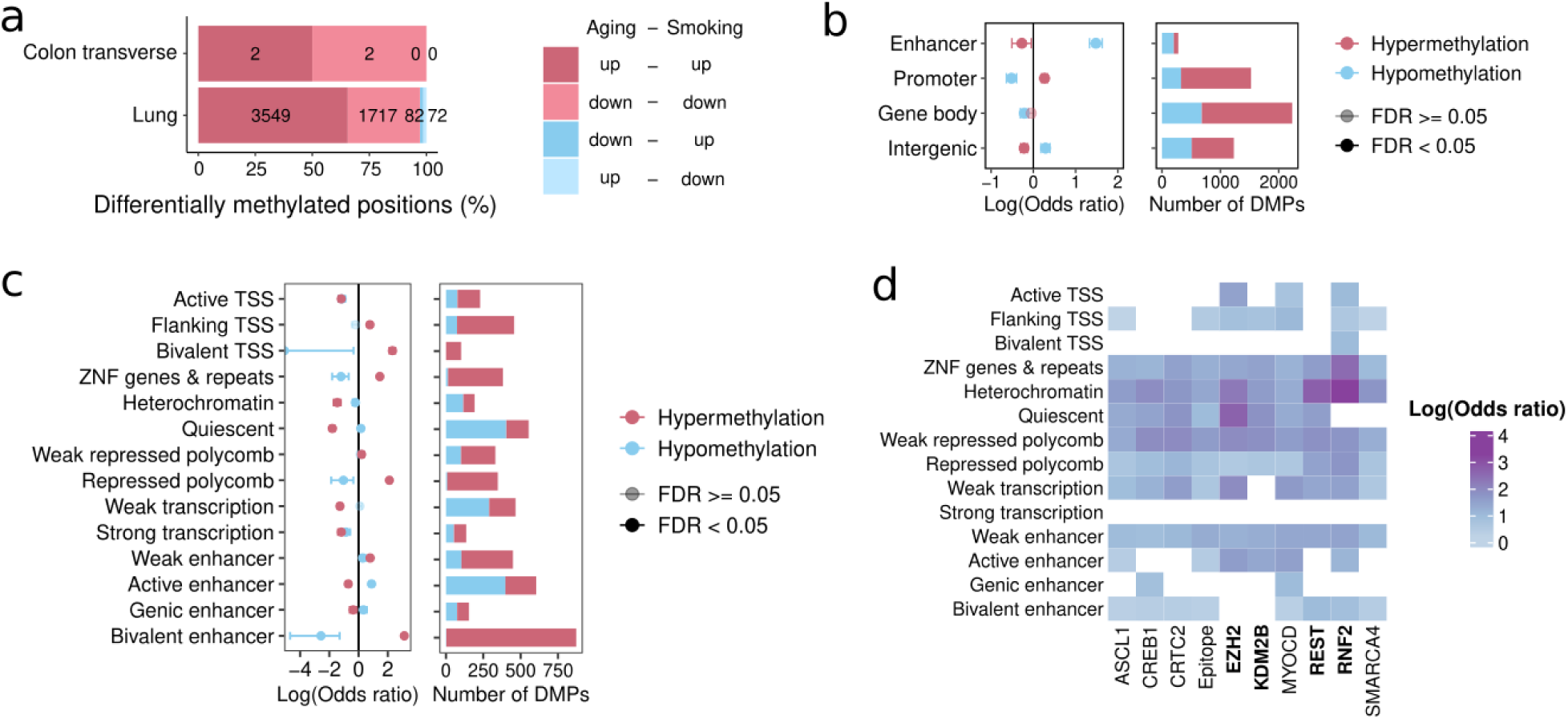
Smoking and age DNA methylation effects. **a**, Tissues with significant concordance in the direction of change (up- or down-regulation) for smoking-age-DMPs (chi-squared tests; FDR<0.05). **b**, Enrichment of smoking-age-DMPs at regulatory regions. **c**, Enrichment of smoking-age-DMPs at chromatin states. **d**, Shared TFBSs enriched in hypermethylation across more than 7 chromatin states for age-smoking-DMPs. TFs highlighted in bold are part of the Polycomb repressive complex.

We then explored the enrichment of age-smoking-DMPs in TFBSs. We find TFs enriched in almost all chromatin states (Fig. S7c). From the most shared TFs in hypermethylated age-smoking-DMPs, we find four components of the Polycomb repressive complex (EZH2, RNF2, KDM2B and REST) enriched in almost all chromatin states (Fig. 6d, Fig. S7d, Table. S11). Hence, both smoking and aging induce similar changes in DNA methylation, particularly through hypermethylation at targets of the Polycomb repressive complex.

A recent study identified CpG sites that are causal for aging-related traits ^61^. We wondered if smoking-induced DNA methylation changes could directly cause tissue aging. We first assessed the degree of overlap between smoking-DMPs with causal aging CpGs. We find that smoking-DMPs significantly overlap causal aging CpGs (Fig. S8A; Fisher’s exact test; FDR<0.05). We then tested if smoking-DMPs were specifically enriched in either protective or damaging DNA methylation changes. Briefly, a protective methylation change is defined as a DNA methylation change that contributes to a positive impact on healthy longevity, while the opposite is true for damaging methylation change^61^. We find that smoking-DMPs are enriched in all four types of aging CpGs (protective hypomethylation, protective hypermethylation, damaging hypomethylation and damaging hypermethylation) with comparable effect sizes (Fig. S8b-c; Fisher’s exact test; FDR<0.05). However, damaging and protective DNA methylation changes have overall negative effects^61^. Thus, our analysis suggests that the impact of cigarette smoking on DNA methylation drives accelerated tissue aging by modifying aging causal CpGs.

### Smoking-induced DNA methylation changes shared with aging are more persistent

Smoking cessation is associated with decreased disease risk and improved life quality ^62,63^. Earlier studies examining gene expression and DNA methylation of the respiratory tract and whole blood in ex-smokers show that while the majority of smoking-induced changes revert after quitting, certain changes persist for years ^22,25,52^. To explore the impact of smoking cessation across human tissues, we performed differential gene expression, alternative splicing and DNA methylation analyses between ex-smokers and smokers, and between ex-smokers and never smokers (Fig. S9a-b). We then classified smoking molecular perturbations as reversible, partially reversible or non-reversible based on whether they change between smokers and ex-smokers, between ex-smokers and never smokers or neither (Fig. 7a-e, Table S13, Table S14). The large majority of genes (92.86%, Fig. 7a), splicing events (96%, Fig. S9c) and CpGs (99.8%, Fig. 7b) were classified as partially reversible. However, in gene expression, there were more reversible genes (6.92%) than non-reversible (0.21%). Conversely, for DNA methylation there were less reversible CpGs (0.05%) compared to non-reversible CpGs (0.14%). Consistent with this, ex-smokers have more partially reversible genes (57%) closer in expression to never smokers (Fig. 7f) but more partially reversible DMPs (52%) closer in methylation levels to smokers (Fig. 7g). Restricting gene expression and DNA methylation analysis to the same donors shows consistent results (Fig. S9d-e). This suggests that ex-smokers are more similar to never smokers in gene expression and to smokers in DNA methylation. To further test this, we trained machine learning classifiers of smokers and never smokers using either gene expression or DNA methylation (Fig. S10a) and used them to reclassify ex-smokers. Gene expression models classify ex-smokers more often as never smokers, with statistically significant differences in lung, artery tibial, thyroid, and adipose subcutaneous (Fig. 7i; binomial test; FDR < 0.05). DNA methylation model classifies 68% of ex-smokers as smokers (Fig. 7i; binomial test; p-value= 0.08). This provides further evidence that ex-smokers are more similar to never smokers in gene expression but more similar to smokers in DNA methylation.

**Fig. 7.**
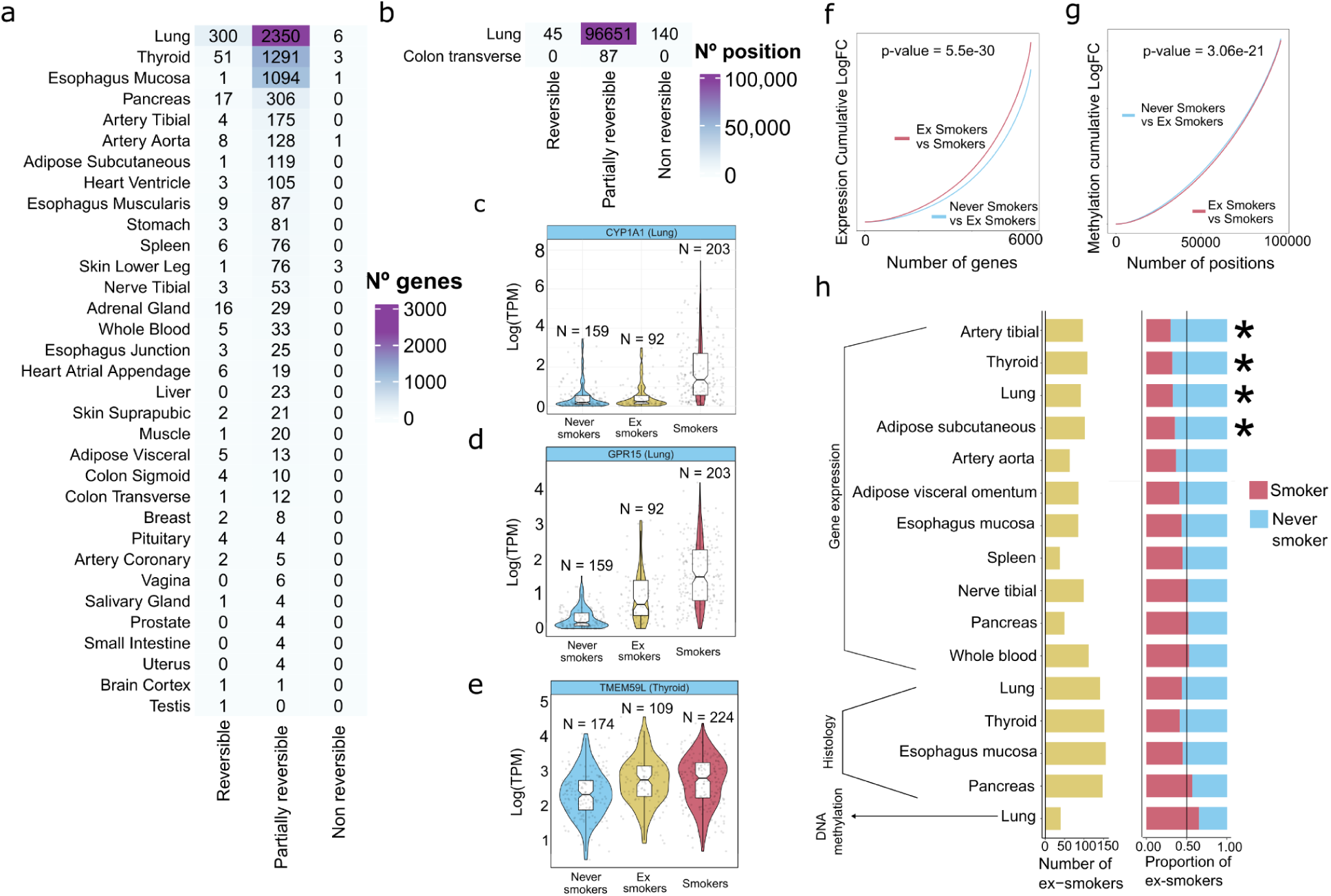
Molecular and histological impact of smoking cessation. **a**, Smoking-DEGs classification into reversible, partially reversible or non-reversible genes per tissue. b, Smoking-DMP classification into reversible, partially reversible and non reversible. **c**, Examples of a reversible, **d**, a partially reversible and **e**, a non reversible gene. **f**, smoking-DEGs log fold change in never vs ex-smokers and in ex-smokers vs smokers. p-value obtained from a Wilcoxon test. **g**, Smoking-DMP log fold change in never vs ex-smokers and in ex-smokers vs smokers. p-value obtained from a Wilcoxon test. **h**, Classification of ex-smoker individuals into smokers or never smokers per tissue in gene expression, histology and methylation. The green bars represent the number of re-classified ex-smokers samples. For gene expression, only highly accurate models (AUC > 0.85) were considered for the classification of ex-smokers.

Finally, we wanted to address whether smoking effects that overlap those of aging were more or less persistent in time compared to those that do not. First, we tested if reversible, partially reversible or non-reversible smoking-DEGs significantly overlap age-DEGs. We find one significant overlap between non-reversible smoking-DEGs and age-DEGs in the skin although the numbers are low (Fisher’s exact test; FDR = 0.0009). In DNA methylation, we find that non-reversible smoking-DMPs in the lung are significantly enriched in age-DMPs (Fisher’s exact test; p-value=1.602e-12), while the reversible and partially reversible smoking-DMPs are not. These results suggest that the smoking effects that affect DNA methylation in common to aging are more persistent in time.

Overall, our results suggest that smoking-induced DNA methylation alterations are more persistent than gene expression changes, especially those that overlap with aging.

## Discussion

Tobacco smoking is the primary cause of preventable mortality ^1^. It has a systemic negative impact on health and increases the risk of developing many diseases ^64^. Here, we provide an exhaustive catalog of molecular changes induced by cigarette smoking throughout the human body and report alterations in many previously unexplored tissues. The effects of smoking on gene expression primarily show tissue-specific effects rather than a shared gene signature, underscoring the importance of a tissue-by-tissue analysis. Nonetheless, we highlight a consistent upregulation of eight genes across multiple tissues. Three of these genes, AHRR, CYP1A1 and CYP1B1, are pivotal components of the aryl hydrocarbon receptor signaling pathway, which is responsible for regulating xenobiotic metabolism ^65^, providing compelling evidence that tobacco toxins reach at least 30 human tissues and systematically activate this pathway to metabolize them. We also report systemic inflammation across tissues that can affect tissue architecture, as we show in the lung regarding macrophage infiltration ^42^. Consistent with this, we find that the smoking-induced DNA methylation changes that correlate with gene expression are located at hypomethylated enhancers of immune-related genes. Furthermore, we report the thyroid as one of the most affected tissues by smoking and characterize its effects on tissue architecture. In particular, we find an enlargement of thyroid follicles, consistent with the association between smoking and an increase in the size of the thyroid gland, named goiter ^34^.

Our findings strongly support previous hypotheses that establish a link between smoking and accelerated aging ^4,5^. We provide evidence that smoking and aging impact gene expression and DNA methylation similarly across tissues. Genes associated with smoking and aging are enriched in immune response across tissues. In addition, aging and smoking drive hypermethylation of bivalent and Polycomb-repressed chromatin states. These regions are transcriptionally inactive regions marked with H3K27me3 ^57^ by the Polycomb-repressive complex. Hypermethylation at Polycomb targets has been previously reported as a systemic effect of aging ^58^ that is evolutionarily conserved ^66^. The reason why epigenetic aging changes accumulate in Polycomb target regions is unclear ^58^. One hypothesis is that systematic DNA damage caused by aging recruits methylation enzymes to sites of DNA repair that will preferentially target CpG-rich regions resulting in hypermethylation in Polycomb-repressed regions ^58,67^. The finding that smoking drives hypermethylation at Polycomb targets as well would be consistent with this hypothesis, as systematic DNA damage also occurs with smoking ^19^. Finally, we show that smoking-driven epigenetic changes are enriched in CpGs causal of aging, suggesting that smoking may in part directly mediate accelerated tissue aging through its effect on DNA methylation.

Overall, our multi-tissue analysis of the impact of cigarette smoking on the human transcriptome, methylome, and histology contributes to a deeper understanding of the physiological and pathological effects associated with smoking, its role as a tissue aging accelerator and the potential health benefits of smoking cessation.

## Methods

### Data collection

All human donors were deceased, with informed consent obtained via next-of-kin consent for the collection and banking of deidentified tissue samples for scientific research. The research protocol was reviewed by Chesapeake Research Review Inc., Roswell Park Cancer Institute’s Office of Research Subject Protection, and the institutional review board of the University of Pennsylvania. For more details on donor characteristics and details on biospecimen collection, sequencing and quality control see the GTEx v8 main paper ^21^.

### Smoking annotation

The smoking annotation was part of the GTEx v8 protected data stored in dbGap (accession number phs000424.v8.p2) and was obtained either from the donors’ medical record or information provided by the donors’ next-of-kin ^68^. We manually curated the annotation and classified donors into never smokers, ex-smokers and smokers. To do so, we used the original MHSMKSTS variable, which encoded information on whether the donor ever smoked and revised the additional comments under MHSMKCMT to distinguish individuals that quit smoking, which were reannotated as ex-smokers. Only cigarette smoking donors were considered. We restricted the analysis to tissues with at least 80 samples with annotation for all the variables included in our models. In total, we analyzed 11,962 RNA-seq samples from 46 human tissues and 717 post-mortem donors.

### Gene and alternative splicing event quantification

Gene and transcript quantifications were based on the GENCODE 26 release annotation (https://www.gencodegenes.org/releases/26.html). We downloaded gene counts and TPM quantifications from the GTEx portal (https://gtexportal.org/home/datasets). We selected genes with the protein-coding and lincRNA biotype on the GTEx GENCODE v26 gtf file. For the expression analysis, we considered expressed genes per tissue (TPM ≥ 0.1 and ≥ 6 reads in ≥ 20% of tissue samples), excluding genes in the pseudoautosomal region (PAR). In total, we analyzed 26,676 genes (19,255 protein-coding and 7,421 lincRNA) across tissues. For the splicing analysis, we downloaded transcripts TPM quantifications from the GTEx portal (https://gtexportal.org/home/datasets) and we used SUPPA2^40^ to calculate percentages of splicing inclusion (PSI) for 7 different types of splicing events: skipped exon, mutually exclusive exons, alternative 3 prime, alternative 5 prime, retained intron, alternative first exon, and alternative last exon. Specifically, we used SUPPA2 to first generate the dictionary of splicing events from the GENCODE v26 annotation and then computed their PSI values for each sample and splicing event. Each splicing event is defined by a set of isoforms: those that include the exonic/intronic sequence (spliced-in isoform) and those that either exclude it or include an alternative exonic sequence (spliced-out isoform). We used the following criteria to select the alternatively spliced events (ASEs) in each tissue: events in protein-coding and lincRNA genes expressed in each tissue; events quantified in all tissue samples; we excluded events with low complexity (fewer than 15 PSI unique values) or insufficient variability (near zero variance); we kept events from expressed isoforms (TPM ≥ 0.5 in ≥ 20% of tissue samples for both the most abundant spliced-in and spliced-out isoforms) and with a quantifiable contribution of the traits of interest (see *Hierarchical partition analysis*). In total, we tested 57,002 alternative splicing events (16,704 skipping exons, 1,142 mutually exclusive exons, 6,491 alternative 5’ splice sites, 7,412 alternative 3’ splice sites, 3,919 intron retentions, 16,681 alternative first exons and 4,653 alternative last exons) across tissues.

### Differential expression analysis

To identify differentially expressed genes (DEGs) we used linear-regression models following the *voom-limma* pipeline ^69,70^. We corrected for technical covariates routinely included in previous GTEx publications ^71,72^. These covariates are related to parameters of donor death (Hardy scale), ischemic time, RNA integrity number (RIN), and sequencing quality control metrics (Exonic rate). We further controlled for unknown sources of variation mainly related to differences in tissue composition and sequencing batch, by including the first two PEER factors ^21,73,74^. We included in the model four demographic traits: the donors’ genetic inferred ancestry ^21^(either European American, African American or admixed donors), sex (either male or female), age, and body mass index (BMI). Ancestry and sex were treated as categorical variables, whereas age and BMI, as continuous variables.

We also corrected for clinical traits obtained from histopathological annotations, publicly accessible from the GTEx portal (https://gtexportal.org/home/histologyPage). However, we only included in the final models the clinical traits with more than 20 diseased individuals associated with changes in gene expression in a simpler model. We first modeled each clinical trait per tissue correcting by the previously mentioned covariates and kept the ones that retrieved at least 5 DEGs (Table S1). Finally, we run one model per tissue correcting by all the technical covariates, the four demographic traits, the chosen clinical traits and smoking.

We compared log-cpm gene expression values and evaluated the statistical significance of smoking and the four demographic traits. Smoking was defined as a categorical variable with three levels: never smokers, ex-smokers and smokers, and all possible contrasts were evaluated. We corrected all analyses for multiple testing using false discovery rate (FDR) through the Benjamini-Hochberg method and considered genes differentially expressed at an adjusted p-value below 0.05.

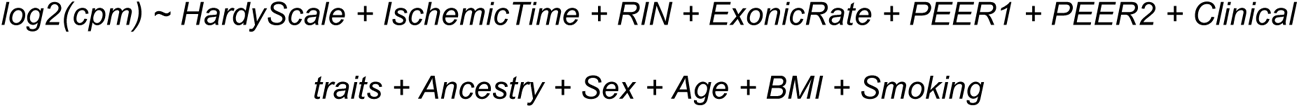

When downsampling we used the same models using an equal number of never smokers and smokers in sets of 50 different permutations. We first performed 50 iterations downsampling to 36 never smokers and 36 smokers, which is the minimum number needed to evaluate all tissues. We then downsampled to 50, 80 and 100 samples per category. To report the results we computed the mean across the 50 permutations.

In order to investigate additive effects we conducted two-tailed Fisher’s exact tests on the common DEGs with smoking and each demographic trait per tissue. The analysis was corrected using FDR through the Benjamini-Hochberg method per demographic trait and considered significant effects with adjusted p-value below 0.05. To check if the direction of change was concordant between demographic traits and smoking, we conducted Chi-square test and corrected for multiple testing in the same way.

To investigate interactions we expanded the linear models in each tissue adding interaction terms between smoking and the other demographic traits (Ancestry:Smoking + Sex:Smoking + Age:Smoking + BMI:Smoking), for the smoking contrast that compares never smokers and smokers. For an interaction to be tested in a tissue we required (1) previously found DEGs with both traits involved in the interaction term, and (2) sufficient sample size, which we define as at least 20 samples per group. To determine the number of samples in each combination we categorized the continuous variables. Age was split into two groups: young (age < 45), and old (age ≤ 45). BMI was divided into three groups: normal (BMI < 25), overweight (25 ≤ BMI < 30) and obese (BMI ≥ 30).

In order to identify the interaction effects between smoking status and genotype on gene expression we followed the steps in ^75^. For each of the gene-variant pairs identified as independent cis-eQTLs in GTEx v8 ^21^, we ran a linear regression model including genotype, smoking status, and covariates to test the effect of interaction term Genotype×Smoking on gene expression separately in each tissue. We tested only the gene-variant pairs that had at least 3 samples in every combination. Each tissue was corrected for a different number of PEER factors based on the tissue sample size, as described in the GTEx eQTL analysis ^21^. For implementation, we used tensorQTL ^76^, where we could model separately for each gene and account for multiple testing errors using Benjamini-Hochberg correction with FDR<0.05.

To validate the genes differentially expressed between smokers and never smokers, we compared our results to other studies that used independent transcriptome datasets. To obtain a list of previously reported smoking-DEGs in lung, we downloaded and parsed Table S1 from Bosse et al. 2012, Supplementary Table S3A and S3B from Landi et al 2008, and Table S1 from Pintarelli et al 2019. For blood, we downloaded and parsed Table S2 from Huan et al 2016 and Table S2 from Vink et al. 2017. For adipose tissue, we downloaded and parsed Table 2 from Tsai et al. 2018. To assess the replicability of our results, we identified smoking-DEGs with GTEx data using similar filtering criteria (logFC and FDR thresholds) as those used in each paper. We performed a two-tailed Fisher’s exact test to test if the number of DEGs we identified significantly overlapped with the genes identified in previous studies. A gene was only considered overlapping if it was DE in the same direction in both studies. We used the protein-coding and lincRNA genes expressed in the respective tissue as background.

### Differential splicing analysis

To perform differential splicing analysis, we used a similar method to the one described in ^47^ to allow both a direct comparison with the differential gene expression analysis and a subsequent quantification of the alternative splicing variation explained by each trait (see *Hierarchical partition analysis*). In short, we modeled Percentage of Spliced In (PSI) values with fractional regressions. This method is suited to work with bounded values that can assume the extremes, as is the case for PSI values in 0 and 1. Specifically, we used the R *glm* function from the R package *stats*^77^ setting *family= ‘quasibinomial* (“logit’’)’ as a parameter. For each splicing event within each tissue, we fitted logit transformed PSI values with the same covariates used in differential expression analysis and evaluated the statistical significance of smoking and the demographic traits of interest: ancestry, sex, age, and BMI.

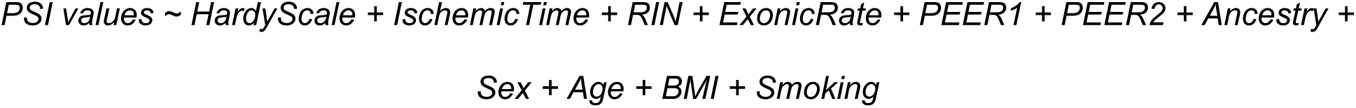

To calculate robust standard errors for our coefficients we used the *vcovHC* function from the R package *sandwich* ^78^ with *type = "HC0"*. To do multiple comparisons for the variables with three levels (e.g., to compute the DSEs between never smokers and smokers, between ex-smokers and smokers, and between never smokers and ex-smokers), we used the function *glht* from the package *multcomp* ^79^, with Tukey’s all-pair comparisons for the *mcp* function. We corrected all analyses for multiple testing using false discovery rate (FDR) through the Benjamini-Hochberg method implemented in the R package stats ^77^. We considered events differentially spliced at an adjusted p-value (FDR) below 0.05.

To investigate the functional consequences of DSEs, we first identified the isoforms that contribute to each splicing event. From these sets of isoforms, we selected the most abundant isoforms per tissue that include (spliced-in) and exclude (spliced-out) the splicing event. Depending on the biotype of these two isoforms, DSEs can then be associated with a switch between protein-coding isoforms, a switch between a protein-coding and a non-coding isoform, or a switch between non-coding isoforms. For those events with a switch between protein coding isoforms, we executed the pipeline from ^47^ to analyze functional consequences of the switches based on functional domains from Pfam ^41^. The percentage of events that belong to each switch type was computed using the sum of the events across tissues from the given switch type over the total number of events, without considering repeated events present due to tissue sharing. To test if smoking tends to be associated with an increase of non-coding isoforms we used a binomial test with a probability of 0.5 as null hypothesis.

### Functional enrichment

We used the R package *clusterProfiler* ^80^ for the different overrepresentation enrichment analyses (ORA) conducted throughout the paper. For each ORA, we defined as background all the expressed genes in the given tissue and analyzed upregulated and downregulated genes independently. We corrected for multiple testing by using the Benjamini-Hochberg method for and considered as significant gene sets with an FDR < 0.05. We performed this analysis across several ontologies, including gene ontology (GO), KEGG and disease ontology (DO). To summarize the enriched GO terms, we employed the *orsumm* package for up- and down-regulated terms in separate ^81^. The analysis was performed by considering all the enriched terms of each tissue at the same time, allowing for an easier comparison between enrichment results.

### Machine learning on histological images

We used the histological images of lung, thyroid, pancreas and esophagus mucosa available in the GTEx Histological Image Viewer (https://gtexportal.org/home/histologyPage), with the aim of building tissue-specific classifiers of smokers and never smokers. For each donor, we downloaded the respective whole slide images (WSI) and divided them into tiles of size 512x512x3 using *PyHist* ^82^ with 2x for the downsample parameter. We kept the tiles with at least 85% tissue content.

We divided the dataset into train and test with the test set being composed of a subset of donors (n = 85) common between the 4 tissues. This step ensures a more robust model comparison. In each tissue, we used 80% of the train subjects for model training, while we kept the remaining 20% for model validation. For modeling we used the pre-trained convolutional neural network (CNN) Xception ^83^ changing only the top layers by adding a Max Pooling ^84^, a Dropout ^84^, and a sigmoid activation ^84^ to get the probability of each subject being a smoker. All models were compiled utilizing a weighted binary cross entropy as loss function, and an *Adam* algorithm with learning rate of 1 × 10^−4^ as an optimizer for the gradient descent. For each tissue we adjusted different class weights in order to mitigate the impact generated by the imbalanced distribution of cases in the final results (see github). The best model for each tissue was selected based on AUC.

### Thyroid follicle detection

We developed a custom CellProfiler ^46^ pipeline to detect and measure thyroid follicles. The first step consists of the computational application of the Ponceau-Fuchsin (PF) stain to the original Hematoxylin and Eosin (H&E) stained image tiles. The PF stain hue absorption helps isolate colloid-containing follicles when using adaptive thresholding in grayscale. We further manually adjusted IdentifyObject module parameters in order to identify the separate follicles. On identified follicles we used the *ObjectMeasurement* module to calculate area, shape and size statistics on all thyroid tiles.

### Cell-type deconvolution

In order to perform cell-type deconvolution of the GTEx bulk dataset we first obtained the processed single cell dataset from GEO GSE173896 ^85^. This dataset integrated 12 individuals (6 smokers, 3 never smokers and 3 ex-smokers) and > 57 000 cells classified into 30 discrete cell types.

Before deconvolution, we processed the single-cell dataset by removing rare cell types (cell types with less than 50 cells across the dataset), and only protein-coding and lincRNA genes in common with the bulk dataset were kept. Next, we used the *cleanup.genes* function from the *BayesPrism* R package to further exclude ribosomal, mitochondrial and sex-chromossome genes, as per package instructions (https://bayesprism.org/pages/tutorial_deconvolution), as well as genes expressed in less than 50 cells. We merged sub cell types into unified cellular populations, and identified marker genes using the *get.exp.stat* function. Marker genes with a log fold change > 0.1 and p.values < 0.01 were selected. Finally, cellular deconvolution was performed based on expression of these genes.

To compare the macrophage populations between smokers groups, while controlling for other sources of variation we fitted a betta regression model on the macrophage proportions in each individual:

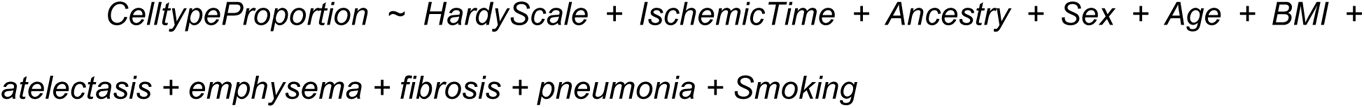

We evaluated the statistical significance of smoking using the *glht* function from the package *multcomp*^86^. We considered the difference significant at a nominal p-value < 0.05

### Differential methylation analysis

We downloaded normalized beta counts of the 754,054 CpGs from the Infinium MethylationEPIC array generated in ^87^. We correlated smoking status with their provided PEERs and found significant correlations with PEER3 and PEER4 (Fig. S4a). Hence, we only corrected for PEER1 and PEER2. We used *limma* ^70^ to run linear models on M values^88^ and corrected for the following set of covariates to be as similar as possible to the previous analysis:

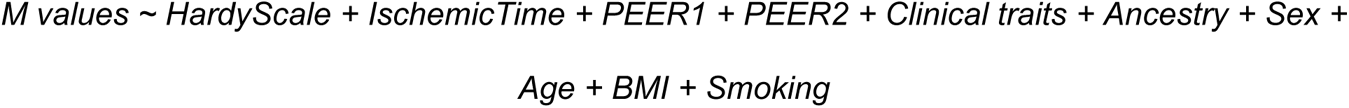

To perform functional enrichment with the EPIC array we need to take into account that some genes contain more probes than others. For this goal, we used the function *gometh* from the *missMethyl* package ^89^ to get GO:BP terms using as background all 754,054 positions studied from the array.

The overlap between smoking and the other demographic traits was followed in the same way as in “*Differential expression analysis”.* We also tested interactions but did not find any significant result.

To validate the DMPs between smokers and never smokers, we compared our results to those obtained with independent datasets using arrays. For lung, we used the results in Table 1 obtained using the 450K array from ^49^, we downloaded the EPIC array data available in Table S2 from ^50^, and the 450K cancer results in Table S1 from ^51^. For blood, we used the data on 450K from ^52^ available in Table S2, the data on EPIC in Table S1 from ^53^, and the data on http://mimeth.pasteur.fr/ from ^60^. For the adipose tissue, we downloaded Table S1 from ^27^ on the 450K array. We performed two-tailed Fisher’s exact tests to check if the DMPs we identified overlapped with the positions observed in previous studies, with the same direction of effect. The background was either the set of studied positions from the EPIC array or the subset of positions available in the EPIC array that were also probed in the 450K array.

We tried to identify interaction effects between smoking status and genotype on DNA methylation following similar steps to section “*Differential expression analysis”,* but we did not find significant results after Benjamini-Hochberg correction with FDR<0.05.

### Annotation of DMPs

The DMPs were classified depending on their location at promoters, enhancers, gene bodies or intergenic based on the annotations provided in the EPIC v1.0 array manifest b5. Positions annotated as “Promoter_Associated” under “Regulatory_Feature_Group” or under “TSS200” or “TSS1500” under “UCSC_RefGene_group” were assigned as promoters. From the rest, the ones with any value in “Phantom5_Enhancers” were assigned as enhancers. From the rest, the positions including “Body”, “1stExon”, “ExonBnd”, “3UTR” or “5UTR” under “UCSC_RefGene_group” were annotated as gene bodies. The rest were annotated as intergenic. We also classified the DMPs into CpG islands, CpG shore, CpG shelf or open sea, using the variable “Relation_to_UCSC_CpG_Island” from the EPIC manifest. To study the enrichment of smoking-DMPs or smoking-age-DMPs in the different genomic locations, we performed Fisher’s exact test for each region separately for hypomethylated and hypermethylated DMPs and adjusted for multiple testing using Benjamini-Hochberg correction.

To annotate the chromatin states around each array probe we used the 18 chromatin states inferred with ChromHMM ^56,57^ for the lung sample BSS01190 generated by the ROADMAP Epigenomics consortium ^57^ and analyzed for EpiMap ^90^. TssFlnkD, TssFlnk and TssFlnkU are reported together under “Flanking TSS’’, EnhA1 and EnhA2, under “Active enhancer” and EnhG1 and EnhG2, under “Genic enhancer”. For the blood analysis we used the PBMC sample BSS01419.

The data was downloaded from https://personal.broadinstitute.org/cboix/epimap/ChromHMM/observed_aux_18_hg19/CALLS/. To perform enrichment of DMPs in the different chromatin states we performed Fisher’s exact test similarly to the analysis on the different genomic locations.

To study the enrichment of transcription factor binding sites around DMPs, we downloaded the processed CHIP-seq data on transcription factors for the human v19 lung available in ChipAtlas^91^. We performed Fisher’s exact test for each transcription factor separately for hypomethylated and hypermethylated DMPs and adjusted for multiple testing using Benjamini-Hochberg correction. The transcription factors part of the Polycomb repressive complex available in the CHIP-seq data were EZH12, SUZ12, YY1, KDM2B, REST, PCGF2, CBX2, CBX8, RNF2, RYBP.

To study the enrichment in causal CpG, we downloaded the causal CpG reported by Ying et al ^61^. For the set of CpG associated with each trait, we computed the significance of the overlap with smoking-CpG and aging-CpG, using Fisher’s exact test. To test the direction of the effect, we classified each causal CpG in protecting or damaging, according to Ying et al, by checking the direction of the product of the Mendelian randomization effect (b) and age effect (B). Sites with b*B > 0 were defined as protective, whereas sites with b*B < 0 were defined as damaging. We conducted an overlap analysis with the smoking-DMPs and aging-DMP using Fisher test. Hyper- and hypomethylated positions were tested independently and p-values were adjusted for multiple testing and

### Correlation between DNA methylation and gene expression

The probe-gene pairs were retrieved first from the EPIC v1.0 manifest b5. We considered a probe to be part of a pair for every gene annotated under “UCSC_RefGene_Name”. For the probes annotated as a promoter or enhancer that did not have any assigned genes in the manifest, we annotated the closest gene using the function *matchGenes* from *bumphunter* ^92^. We computed Pearson’s correlation for the DMP-DEG pairs and corrected for multiple testing using the Benjamini-Hochberg method. We considered significant correlations with an FDR < 0.05. To compute the percentage of DEGs that significantly correlate with DMPs, we considered DEGs associated with at least one probe in the array.

### Classification into different reversibility categories

In order to test the reversibility of gene expression, alternative splicing events and DNA methylation, we compared the lists of DSGs/DSEs/DMP between groups: never smokers, ex-smoker and smokers and classified them into 3 categories.

● Reversible when the smoking-DEGs/DSEs/DMP (never smoker vs smokers) were also found with the same direction in ex-smokers vs smokers but not in ex-smoker vs smokers;
● Non reversible, for smoking-DEGs/DSEs/DMP found also in the never smokers vs ex-smokers comparison in the same direction but not in the ex-smokers vs smokers analysis;
● Partially reversible genes, when the events were not classified as either reversible or non reversible.

For partially reversible genes, we further compared the mean absolute logFC obtained in the never smokers vs ex-smokers analysis with the one obtained in the ex-smokers vs smokers analysis through a two-tailed Wilcoxon paired test.

### Machine learning on gene expression and methylation data

All the machine learning models based on gene expression and methylation were trained using the gradient boosting algorithm implemented in the *lgb.train* function from the *lightgbm* R package.

For gene expression, values were normalized in log2 scale, log2(TPM +1) and only protein-coding and lincRNA genes were considered as input. For methylation data all the probes in lung data were considered. In order to evaluate model performance, a 5-fold cross validation scheme was implemented.

Since the models had good accuracy (cross-validated mean AUC > 0.85), we trained a final model using all samples from smokers and never smokers for each tissue. Feature importance was based on this final model and was obtained with the *lgb.plot.importance* function, setting the measure parameter to “Gain”. Classification of ex-smokers samples into never smokers or smokers was performed using the final model.

## DATA AVAILABILITY

All GTEx protected data are available at dbGap under the accession number phs000424.v8. (https://gtexportal.org/home/protectedDataAccess). Expression and histology data is publicly available through the GTEx Portal as downloadable files (https://www.gtexportal.org). DNA methylation data is available in GEO under the accession number <u>GSE213478</u>. The single-cell data is available in GEO under the accession number GSE173896.

Analysis scripts are available at https://github.com/Mele-Lab/2024_GTEx_Smoking.

## ACKNOWLEDGMENTS

J.M.R. was supported by a predoctoral fellowship from “la Caixa” Foundation (ID 100010434) with code LCF/BQ/DR22/11950022. R.R was supported by the scholarship BD/07092/2021 from Fundação para a Ciência e a Tecnologia (FCT) and the European Social Fund. P.G.F. acknowledges the grant 2022.15770.CPCA.A1 from RNCA-FCT. M.M. was supported by a grant PID2019-107937GA-I00 funded by MCIN/AEI/10.13039/501100011033 and a grant RYC-2017-22249 funded by MCIN/AEI/10.13039/501100011033 and by “ESF Investing in your future”. We thank Kristin Ardlie (Broad Institute, USA), François Aguet (Illumina Inc, USA) and Roderic Guigó (CRG, Spain) for useful discussions at the start of this work and Winona Oliveros (BSC, Spain) for discussions at the end of the work. We thank Daniel Martinez and Jose Ramirez from Hospital Clinic (UB, Spain) for useful insights on visual identification of smoking-borne histological changes. Fig. 1 was created with <u>BioRender.com</u>.

## Declaration of interests

The authors declare no competing interests.

## List of supplementary tables

Table S1 - Clinical traits included per tissue

Table S2 - Smoking-DEGs per tissue (including reversible status)

Table S3 - Enrichment analysis of all terms (per tissue) and all Ontologies (GO:BP + KEGG + DO for thyroid and Lung)

Table S4 - Enrichment analysis - Orsum

Table S5 - Smoking-DSEs per tissue. It includes a column with the most expressed isoform including and excluding the event and the protein domains where the event is involved for those isoforms

Table S6 - Estimated cellular proportion of macrophages in the lung

Table S7 - Features measured on thyroid follicles identified on thyroid histopathological images via CellProfiler

Table S8 - Functional enrichments of age-smoking DEGs

Table S9 - DMP in each tissue.

Table S10 - Functional enrichment per chromatin region of smoking-DMPs, age-smoking-DMPs, smoking-DMPs without DMPs sitting in TFBSs from Polycomb proteins, and age-smoking-DMPs without DMPs sitting in TFBSs from Polycomb proteins.

Table S11 - TFBSs enrichment of DMPs.

Table S12 - Functional enrichment of smoking-DEGs correlated with smoking-DMPs

Table S13 - Never vs ex and Ex vs Smoker analysis

Table S14 - Never vs ex and Ex vs smokers DMP analysis

## Supplementary figures

**Fig. S1.**
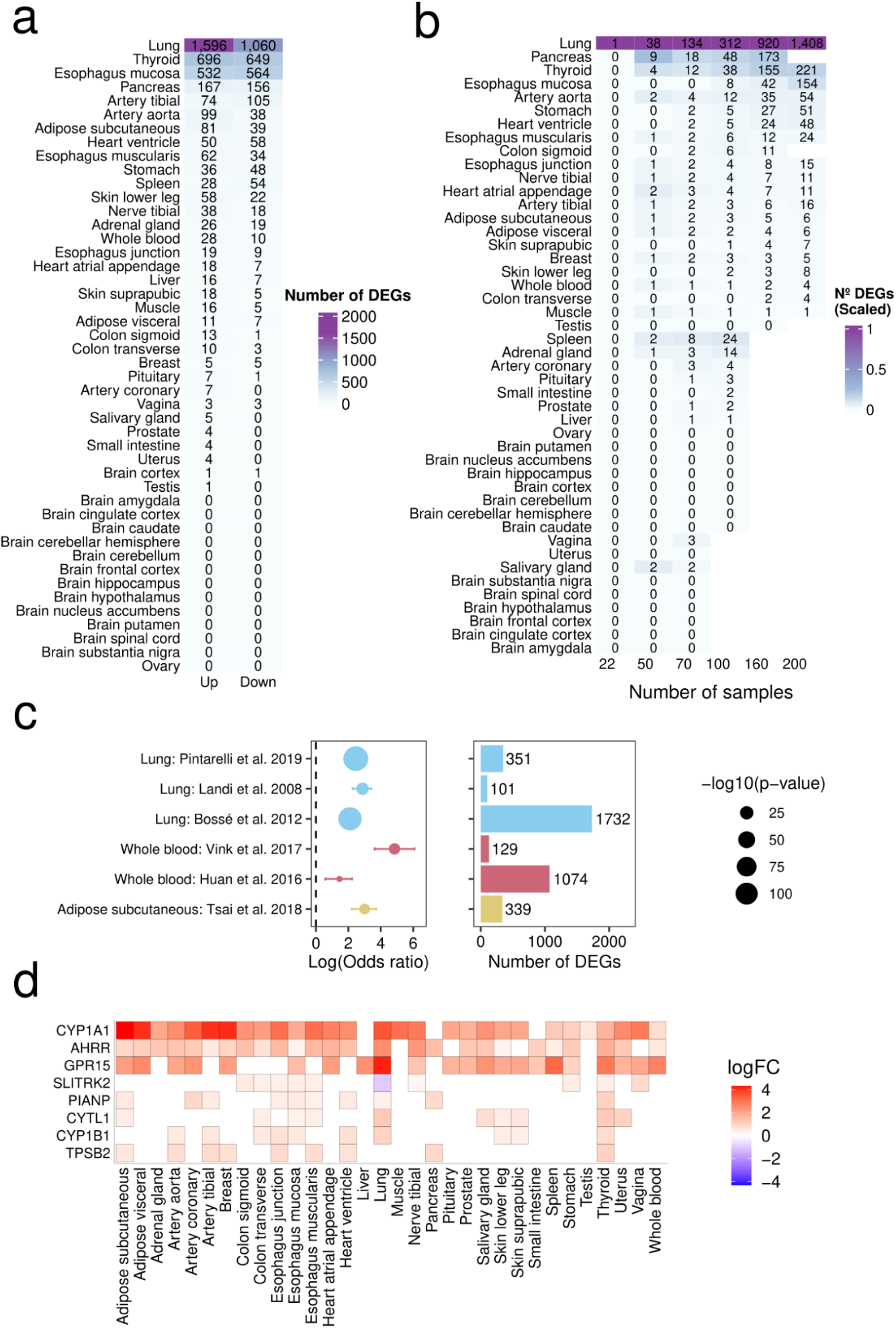
Differential expression analysis. **a**, Number of upregulated and downregulated smoking-DEGs per tissue. **b**, Median number of smoking-DEGs per tissue when downsampling in 50 permutations. The x axis represents the different sample sizes, divided into equal numbers of smokers and never smokers. **c**, Replication of the smoking-DEGs found in this work with previous literature. The dot plots show the log odds ratio from the two-sided Wilcoxon-test, with respective confidence intervals, while the bar plot shows the number of DEGs in each study. Statistical significant overlaps were found with all studies. **d**, Log fold changes of the recurrent genes across tissues. A white square represents a tissue where that gene was not found differential expressed (FDR > 0.05).

**Fig. S2.**
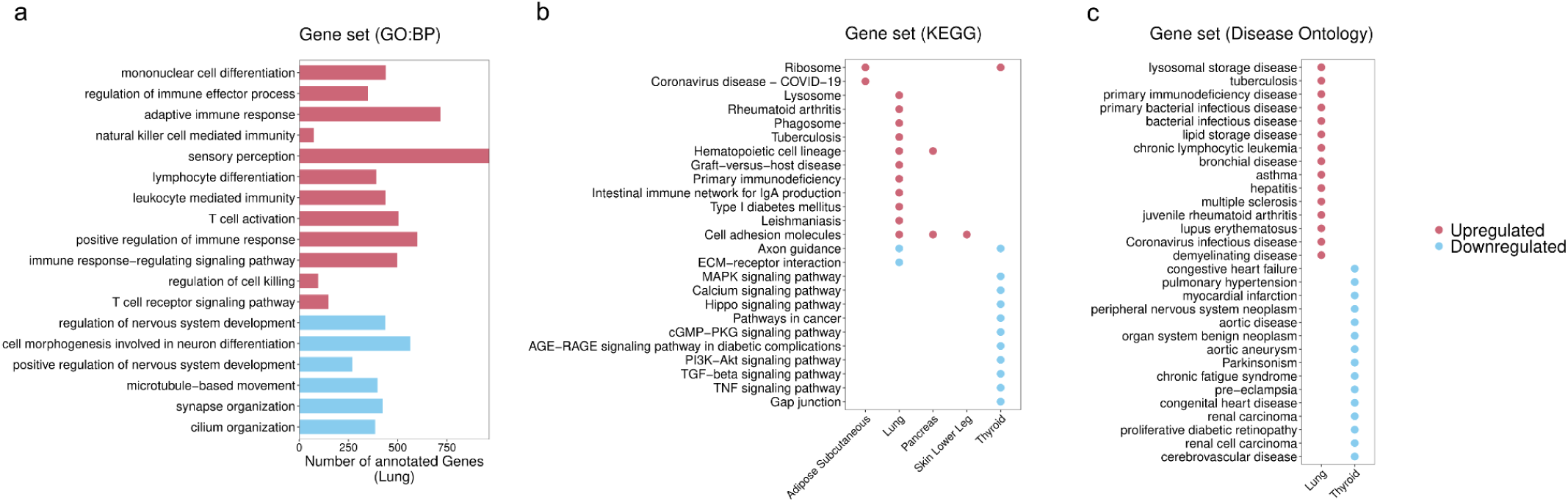
Enrichment analysis performed on smoking-DEGs across tissues. **a**, Top enriched gene ontology (biological process) in lung for upregulated (red) and downregulated (blue) genes in lung (FDR < 0.05 and minimum gene count > 5). **b**, Top enriched KEGG terms (FDR < 0.05 and minimum gene count > 5), for upregulated genes (red) and downregulated genes (blue) across tissues. Only the top 15 terms per tissue are represented. **c**, Top enriched disease ontology terms enriched in lung and thyroid (FDR < 0.05 and minimum gene count > 5).

**Figure S3.**
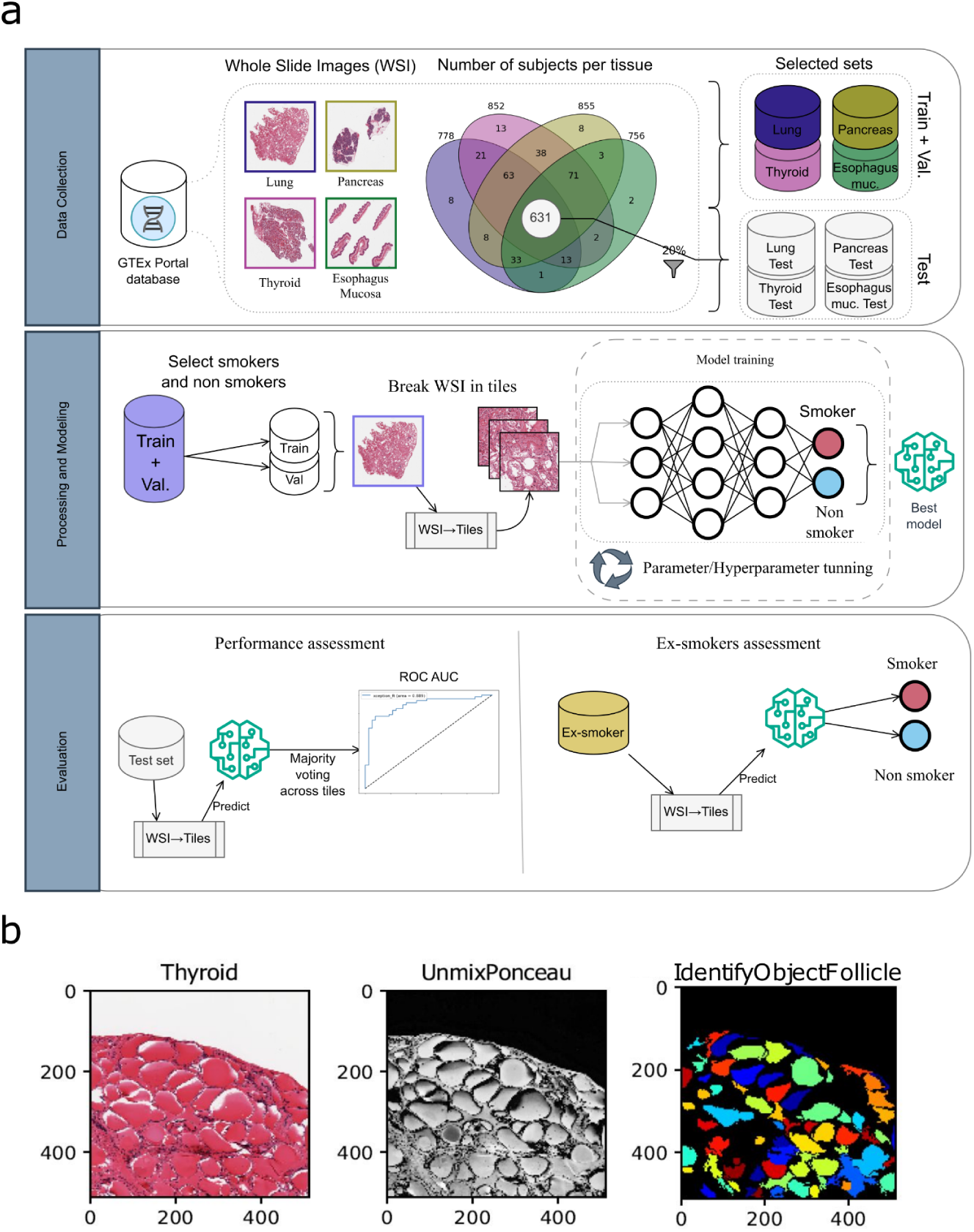
Histology Images analysis. **a**, General scheme of the machine learning approach used in this work for histology analysis. Whole slide images (WSI) for lung, thyroid, pancreas and esophagus mucosa were downloaded from the GTEx portal database. We split the data into train, validation (val) and test set. The test set was derived by taking 20% (126 subject) of the common samples across all tissues (631). The remaining samples from each tissue were further divided into train (80%) and validation (20%). For model training, WSI were divided into tiles, and kept tiles with at least 85% tissue content. For modeling, we used the pre-trained convolutional neural network (CNN) Xception changing only the top layers. Since the model outputs tile specific prediction, we used a majority voting approach to determine subject level prediction. The final model performance was evaluated in the test set. Finally, these models were used on ex-smoker samples to be classified as smoker or never smoker. **b**, CellProfiler pipeline illustration of a tile. Left — the original H&E stained tile of a thyroid, Center — Grayscale tile after computational transform with the Ponceau-Fuchsin stain, Right — Identified individual thyroid follicles (color).

**Fig. S4.**
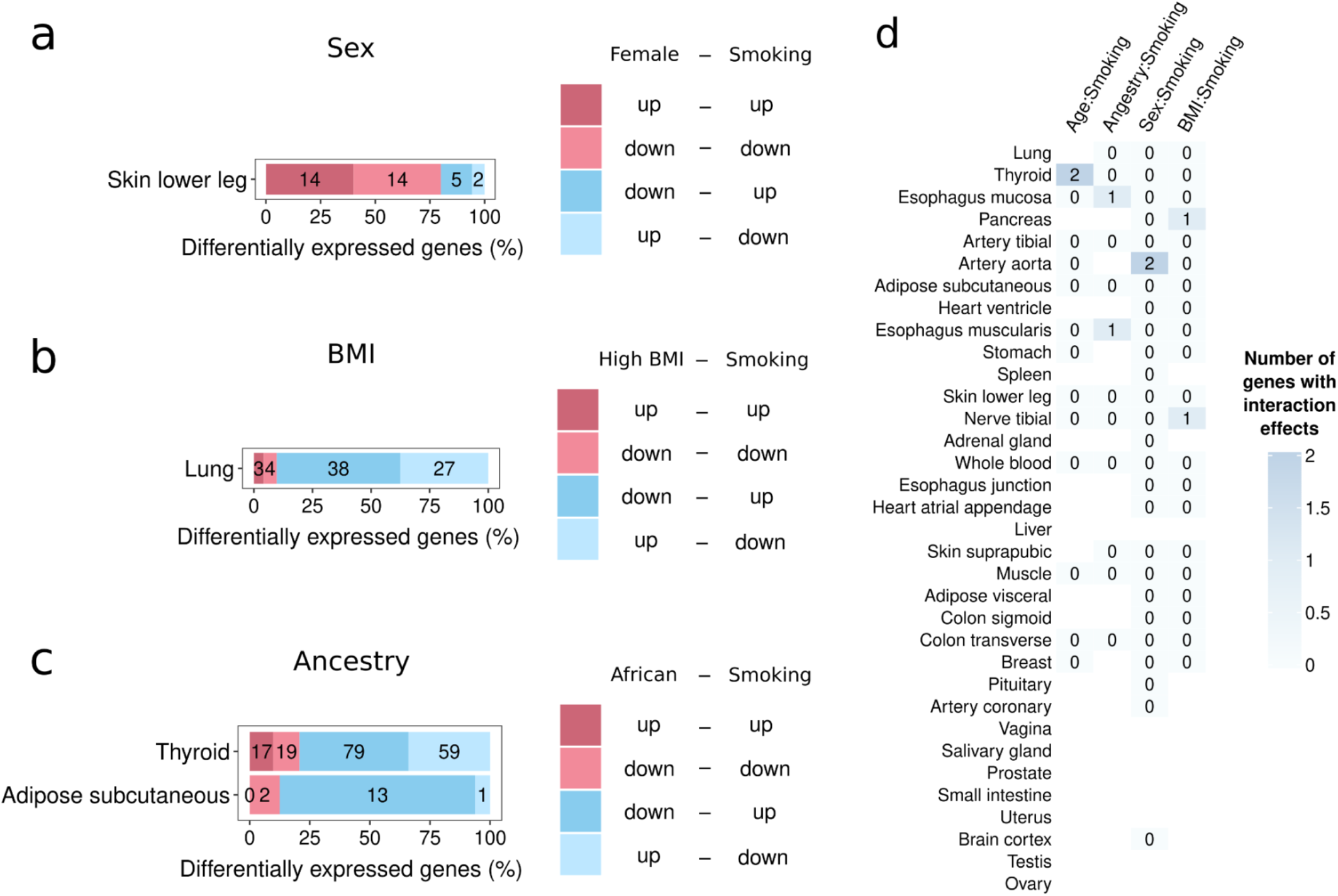
Additive and interaction effects. **a-c**, Bar plots showing the tissues with significant concordance in the direction of change (up- or down-regulation) for smoking-sex-DEGs, smoking-BMI-DEGs and smoking-ancestry-DEGs across tissues (chi-squared tests FDR<0.05). **d**, Genes with interaction effects between smoking and the different demographic traits.

**Fig. S5:**
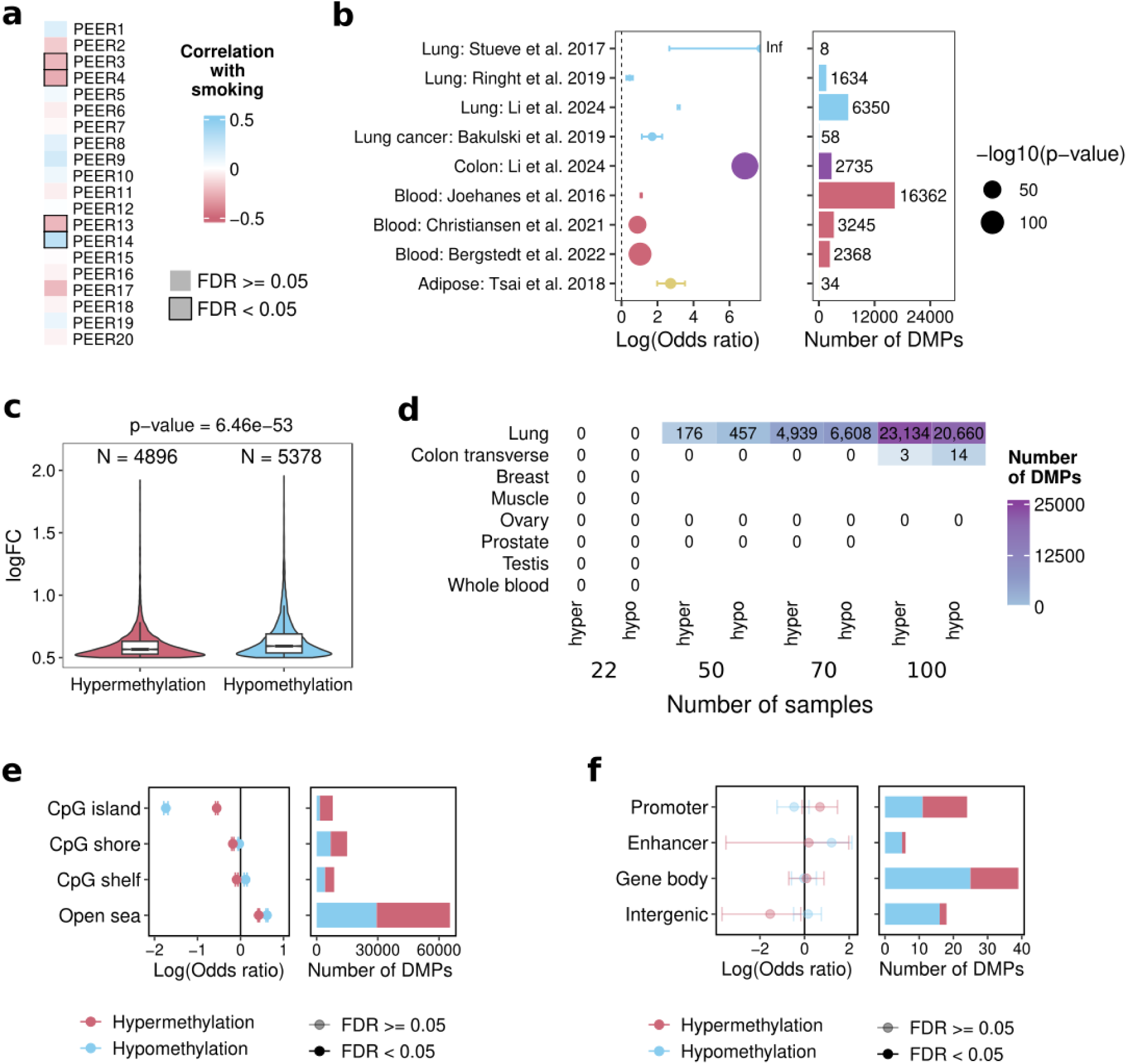
Association between smoking and DNA methylation across tissues. **a**, Correlation between smoking and PEER factors. **b**, Replication of smoking-DMPs with DMPs identified in previous studies (Fisher’s exact tests<0.05). The overlap in colon was performed with our colon DMPs, whereas all the rest were performed with our lung DMPs. Y-axis shows the article reference and the tissue used in the respective study. Left plot dashed line indicates odds ratio = 1, values above are enriched and below are depleted. Colors refer to the tissue used in the studies. Right plot shows the number of DMPs reported in each study that we can test. **c**, Effect sizes of hypermethylated and hypomethylated positions in lung. **d**, Number of DMPs obtained when downsampling to different number of samples (x-axis). The number of smokers and never smokers is the same. The cell numbers correspond to the mean of 50 different permutations per tissue and number of samples. **e**, Enrichment of smoking-DMPs on EPIC annotations about CpG context. **f**, Enrichment of colon transverse smoking-DMPs at regulatory regions.

**Fig. S6:**
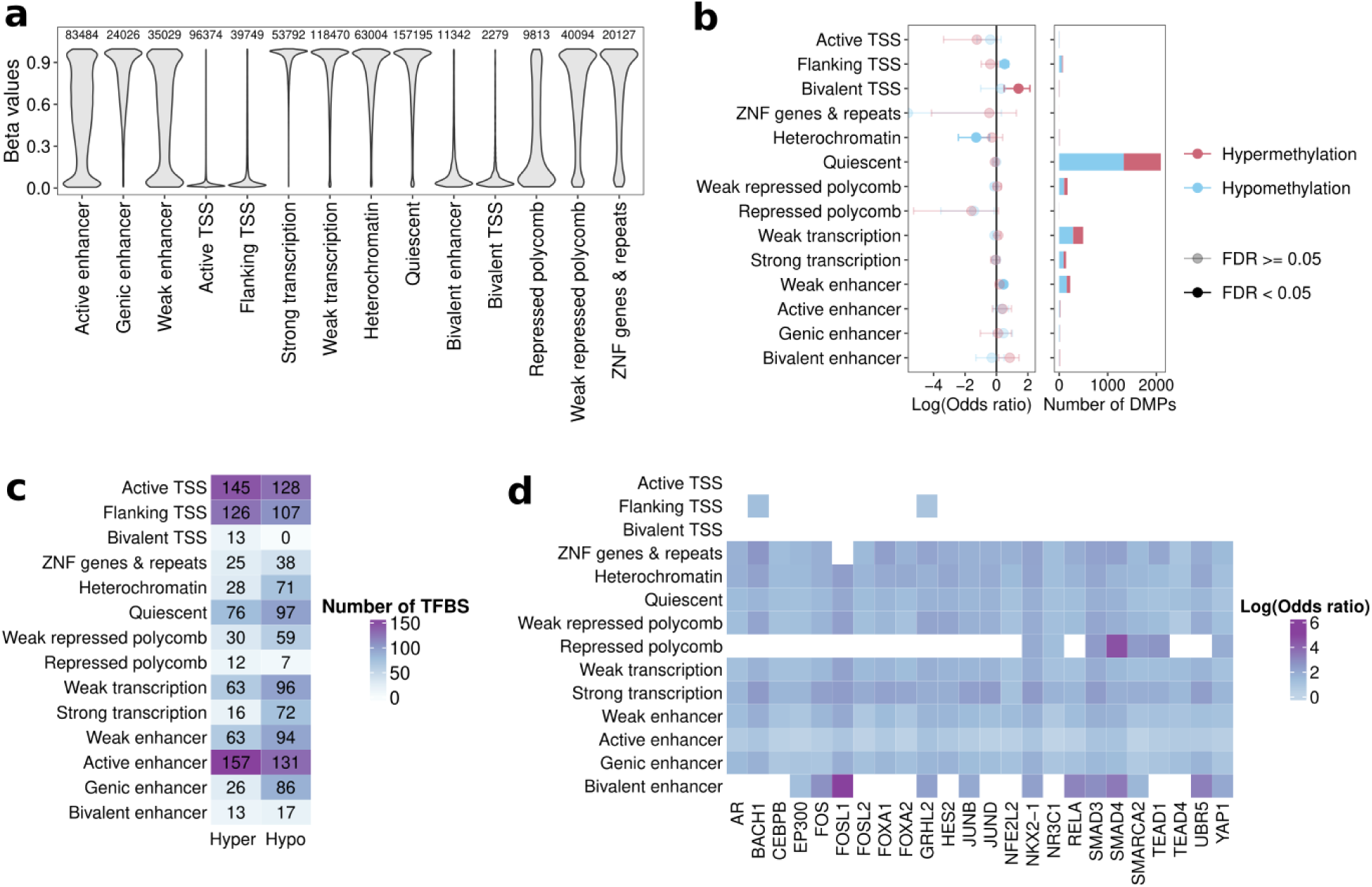
Association between smoking and DNA methylation across chromatin states. **a**, Methylation levels in never smokers stratified by chromatin state. **b**, Enrichment of smoking-DMPs from Christiansen et al. at blood chromatin states **c**, Number of TFBSs enriched in hypermethylated and hypomethylated CpGs per chromatin state **d**, Shared TFBSs enriched in hypomethylation across more than 8 chromatin states.

**Fig. S7.**
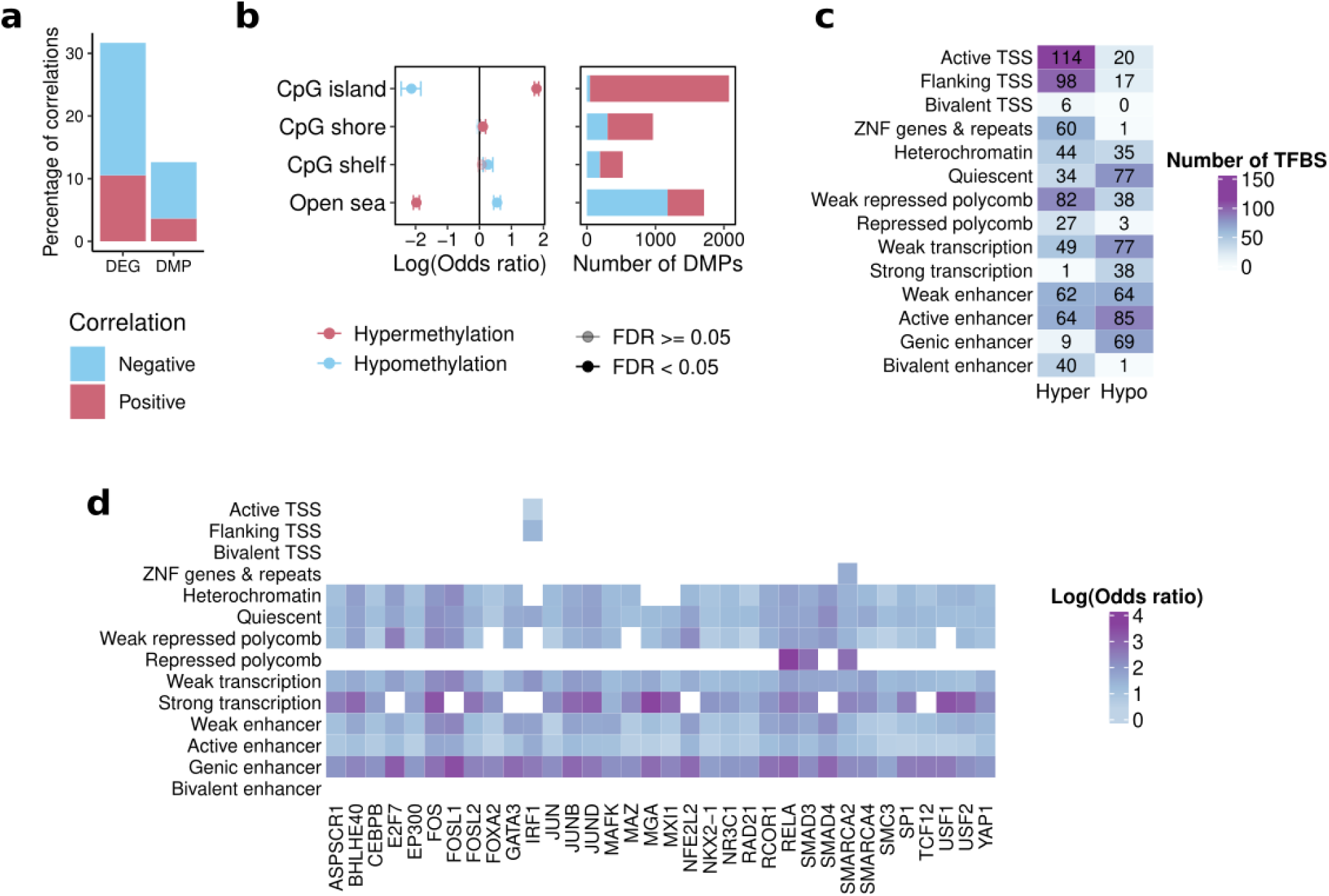
Correlation between DNA methylation and expression and enrichments of smoking-age-DMPs. **a**, Percentage of DEG-DMP significantly correlated over the total number of DEGs or DMPs tested. **b**, Enrichment of smoking-age-DMPs depending on CpG context. **c**, Number of TFBSs enriched in hypermethylated and hypomethylated CpGs per chromatin state for age-smoking-DMPs **d**, Shared TFBSs enriched in hypomethylation across more than 6 chromatin states for age-smoking-DMPs.

**Fig. S8.**
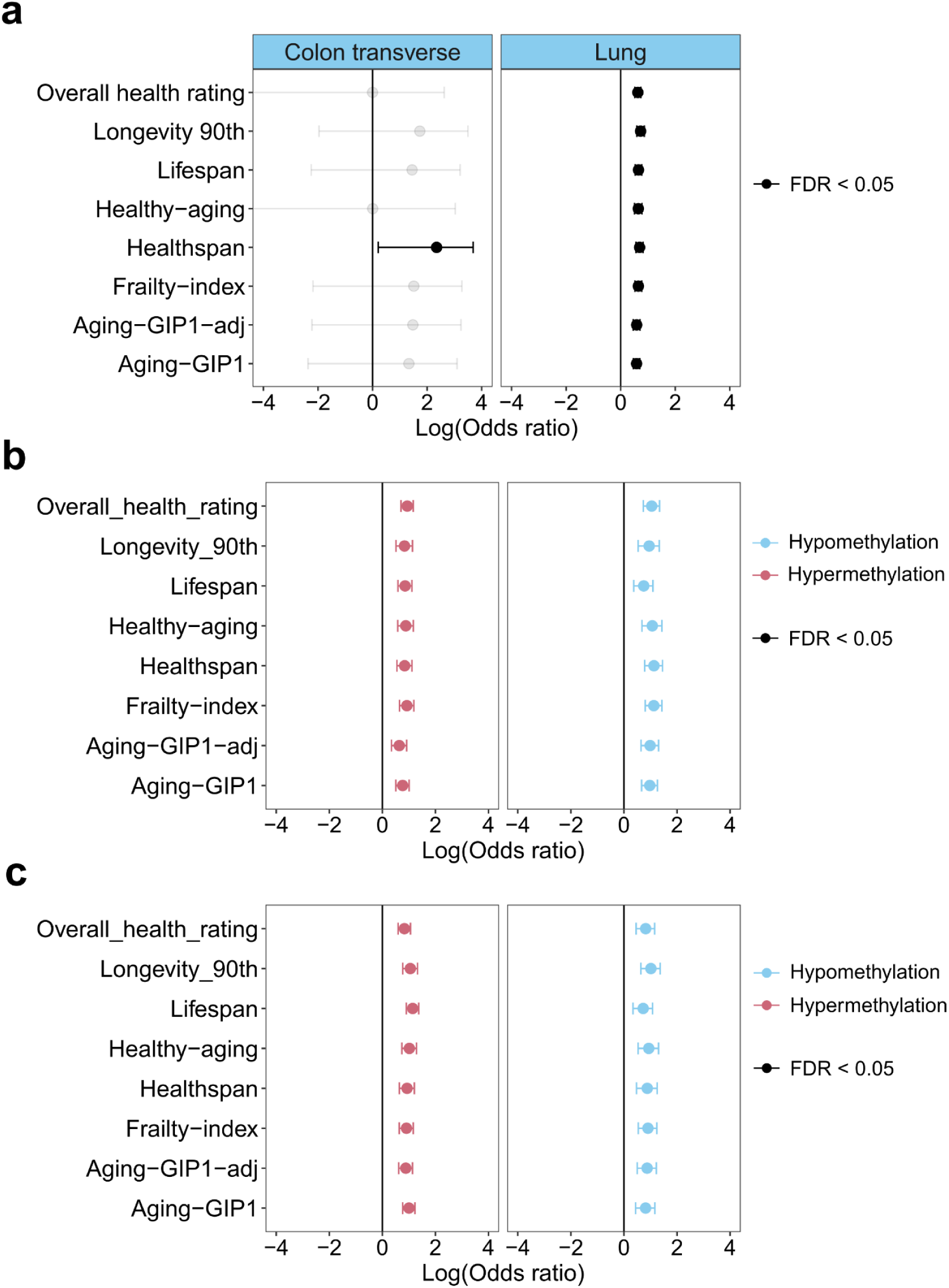
Overlap between smoking-DMPs and causal aging CpG. **a**, Overlap between smoking-DMP in colon transverse (left) and lung (right) with causal aging CpG position from Ying et al. (63). **b-c,** Overlap between smoking-DMPs in lung with damaging (**b**) and protective (**c**) CpG positions, discriminated by hypermethylated positions (in red) and hypomethylation positions (in blue).

**Fig. S9.**
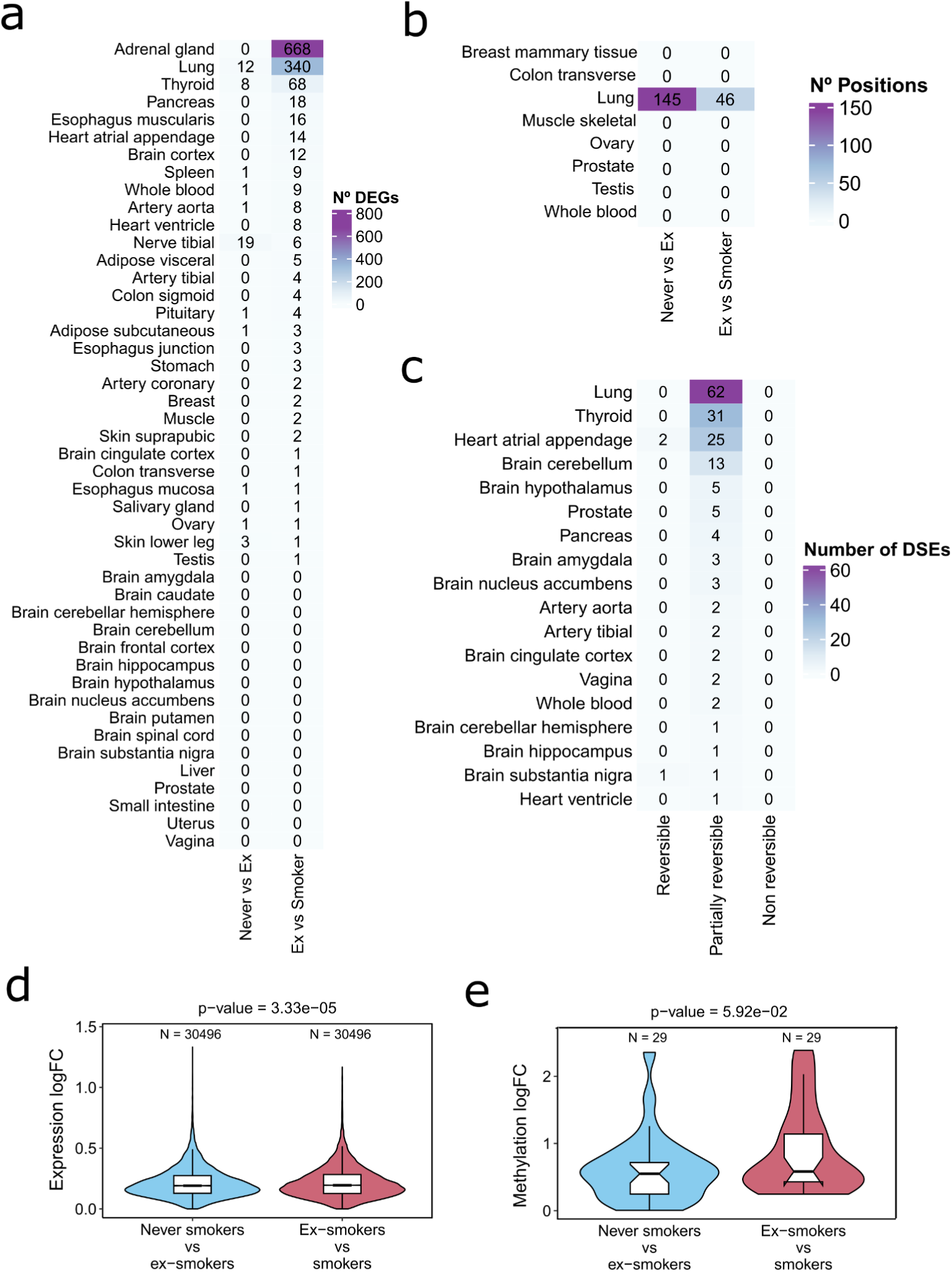
Reversibility analysis. **a**, Number of differentially expressed genes between never smokers and ex-smokers, and ex-smokers and smokers. **b**, Number of differentially expressed positions between never smokers and ex-smokers, and ex-smokers and smokers. **c**, Number of reversible and non reversible splicing events. **d,** smoking-DEGs and **e**, smoking-DMP log fold change in never vs ex-smokers and in ex-smokers vs smokers when analysis was constrained to the common samples between methylation and gene expression.

**Fig. S10.**
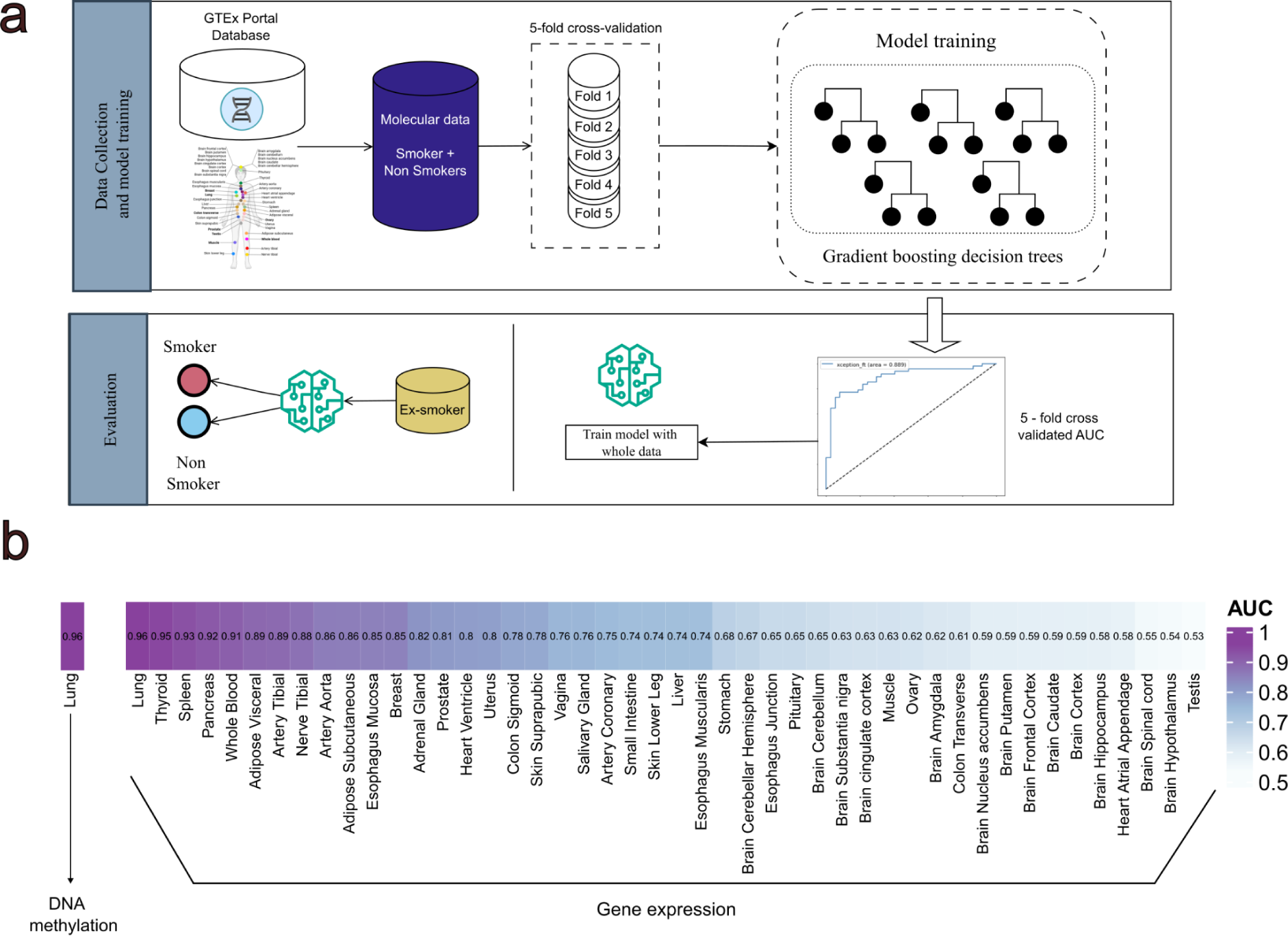
Machine learning to reclassify ex-smokers. **a**, General scheme of the machine learning approach used in this work for gene expression and methylation. Briefly, never smokers and smokers samples were considered as the target classes for model training. Performance was assessed using a 5-fold cross-validation scheme. After assessment, the binary classification model was used to predict in which of these classes (smokers and never-smokers) the ex-smokers samples were classified. **b**, AUC across tissues (average across 5-fold cross-validation). **c**, Feature importance across tissues. The numbers in the heatmap represent the ranking of the importance of each gene for the machine learning model. Only the top 3 genes for each tissue are represented. **d**, Top 10 most important features for the machine learning classifier to distinguish smokers and never smokers.

## Notes

### Competing Interest Statement

The authors have declared no competing interest.

### Summary of Updates

We improved some sections and changed the title of the manuscript

